# How many bird and mammal extinctions has recent conservation action prevented?

**DOI:** 10.1101/2020.02.11.943902

**Authors:** Friederike C. Bolam, Louise Mair, Marco Angelico, Thomas M. Brooks, Mark Burgman, Claudia Hermes, Michael Hoffmann, Rob W. Martin, Philip J.K. McGowan, Ana S.L. Rodrigues, Carlo Rondinini, Hannah Wheatley, Yuliana Bedolla-Guzmán, Javier Calzada, Matthew F. Child, Peter A. Cranswick, Christopher R. Dickman, Birgit Fessl, Diana O. Fisher, Stephen T. Garnett, Jim J. Groombridge, Christopher N. Johnson, Rosalind J. Kennerley, Sarah R.B. King, John F. Lamoreux, Alexander C. Lees, Luc Lens, Simon P. Mahood, David P. Mallon, Erik Meijaard, Federico Méndez-Sánchez, Alexandre Reis Percequillo, Tracey J. Regan, Luis Miguel Renjifo, Malin C. Rivers, Nicolette S. Roach, Lizanne Roxburgh, Roger J. Safford, Paul Salaman, Tom Squires, Ella Vázquez-Domínguez, Piero Visconti, James R.S. Westrip, John C.Z. Woinarski, Richard P. Young, Stuart H.M. Butchart

## Abstract

Aichi Target 12 of the Convention on Biological Diversity (CBD) aims to ‘prevent extinctions of known threatened species’. To measure its success, we used a Delphi expert elicitation method to estimate the number of bird and mammal species whose extinctions were prevented by conservation action in 1993 - 2020 (the lifetime of the CBD) and 2010 - 2020 (the timing of Aichi Target 12). We found that conservation prevented 21–32 bird and 7–16 mammal extinctions since 1993, and 9–18 bird and 2–7 mammal extinctions since 2010. Many remain highly threatened, and may still become extinct in the near future. Nonetheless, given that ten bird and five mammal species did go extinct (or are strongly suspected to) since 1993, extinction rates would have been 2.9–4.2 times greater without conservation action. While policy commitments have fostered significant conservation achievements, future biodiversity action needs to be scaled up to avert additional extinctions.

## Introduction

The Parties to the Convention on Biological Diversity (CBD) adopted an ambitious strategic plan for the decade 2011-2020, comprising 20 ‘Aichi Biodiversity Targets’. Target 12 states that ‘*By 2020, the extinction of known threatened species has been prevented and their conservation status, particularly of those most in decline, has been improved and sustained*’. A mid-term assessment concluded that further extinctions were likely by 2020, but that conservation measures had prevented some extinctions, with further prevented extinctions likely before 2020 (CBD 2014).

Considering compelling evidence of a continued deterioration of the state of nature in the face of increasing pressures (IPBES 2019, Díaz et al., 2019), investigating the impact of conservation efforts is key to evaluating whether we have the knowledge and techniques to reverse negative trends, and to galvanise further action. Previous assessments of conservation impact investigated whether trends in extinction risk as measured through the Red List Index would have changed if no species had improved in conservation status (Hoffmann et al., 2010, Szabo et al., 2012), or if no conservation actions had taken place (e.g. Hoffmann et al., 2015, Young et al., 2014). Butchart et al. (2006) estimated which bird species would have gone extinct without conservation during 1994-2004, based on expert knowledge. Looking ahead, green listing will provide standardised methods to quantify the impact of conservation compared with a counterfactual (Akçakaya et al., 2018).

Here we build on these studies to quantify the extent to which ‘*the extinction of known threatened species has been prevented*’ by conservation action, by identifying those bird and mammal species that would have gone extinct without such action during 1993-2020 (the lifetime of the CBD) and 2010-2020 (approximately the timing of Aichi Target 12). We focused on birds and mammals as some of the best documented taxonomic Classes on the IUCN Red List, and with disproportionate attention from the public (Troudet et al., 2017) and for recovery planning (Walsh et al., 2013). We did this for each time period by: (a) identifying candidate species that could plausibly have gone extinct without conservation action; (b) documenting for each species the key factors necessary to evaluate whether the actions implemented could plausibly have prevented its extinction; and (c) using a Delphi expert elicitation approach (Hemming et al., 2018; Mukherjee et al., 2015) to estimate the probability that each candidate species would have gone extinct in a counterfactual scenario where no conservation action would have taken place. We then combined our results with the number of known extinctions to quantify the effect of conservation on observed extinction rates.

## Methods

### Identifying candidate species

We first identified bird and mammal species that were listed on the IUCN Red List (IUCN 2019) as Extinct in the Wild, Critically Endangered under any criteria, or Endangered under Criterion D (i.e. with a population size below 250 mature individuals) at any point since 1993. This initial list included 368 bird and 263 mammal species. Two groups of 2-4 authors reviewed independently whether each species was a plausible candidate, then we compared results, discussed discrepancies, and agreed a shortlist of candidate species comprising 48 bird and 25 mammal species.

For each candidate species we compiled standardised information from the IUCN Red List on their population size and trends in 1993, 2010 and in the latest assessment year, as well as on threats and conservation actions, and summarised the key arguments to assess the probability if they may have gone extinct without conservation. Further review of the species information by key taxon experts reduced the final candidate list to 39 bird and 21 mammal species. We considered a shorter list of 23 bird and 17 mammal species for 2010-2020, by excluding species whose populations in 2010 were sufficiently large that extinction by 2020 was deemed implausible.

### Delphi process

We used a Delphi expert elicitation method following Hemming et al. (2018) and Mukherjee et al. (2015). We asked 28 bird and 26 mammal evaluators to estimate independently the probability that each candidate species would have gone extinct without conservation action. Specifically, we asked evaluators three questions for each time period: *Realistically, what do you think is the (1) lowest plausible probability / (2) highest plausible probability / (3) best estimate for the probability that conservation action prevented extinction for this species during the period (i*.*e. what is the probability that, if action had ceased in 1993/2010, and no subsequent actions were implemented, the species would have gone extinct by 2020)?*

To answer these questions, evaluators were instructed to use the information summarised for each species as well as any other information they had access to, and to assume that all conservation action would cease at the start of the period, for example hunting bans overturned, protected status revoked, protected areas degazetted, and captive breeding programmes ceased.

For each species, and for each time period, we calculated the median lowest (question 1), highest (question 2) and best estimate (question 3) of probabilities that extinction has been prevented (von der Gracht, 2012), and level of agreement (see supplementary material). The median scores for all species were shared with all evaluators, followed by teleconference calls in which evaluators discussed each species in turn. Evaluators were then given the opportunity to revise their scores (again independently and anonymously) to incorporate any insights gained during the call.

### Analysis

We summarised the overall results as the number of species whose extinction had been prevented as X–Y, with X representing species with a median best estimate ≥90% that extinction had been prevented and Y representing species with a median best estimate >50%, following an analogous approach for defining Extinct and Critically Endangered (Possibly Extinct) species (Butchart et al., 2018). We treated species assessed as Extinct in the Wild as having 100% probability that their extinction was prevented, because they would be Extinct if not for their captive populations.

For all species with a median best estimate >50% for 1993-2020, we analysed their distribution, threats, actions implemented, current IUCN Red List category, and current population trend (see Figures S1 - S7 for plots of all candidates and for species with a median best estimate >50% for 2010-2020). Finally, we compared the total number of those species with numbers of species confirmed or strongly suspected to have gone Extinct in the same period according to the IUCN Red List (Tables S2, S3).

## Results

Of 39 candidate bird species for the 1993-2020 period, 15 had a median best estimate ≥90% that their extinction was prevented (Fig. 1a), with a further 11 species having a median best estimate >50%. Agreement among evaluators for these 26 species was high for 14 and medium for 12 species. Including six additional species listed as Extinct in the Wild during the time period (Table S4), we consider that 21–32 bird species would have gone extinct without conservation during 1993-2020. In contrast, there were 10 confirmed or suspected extinctions since 1993 (Table S2). Hence, in the absence of conservation, the total number of bird extinctions since 1993 would have been 3.1–4.2 times higher (31–42 vs. 10) (Table S3).

**Figure 1.**
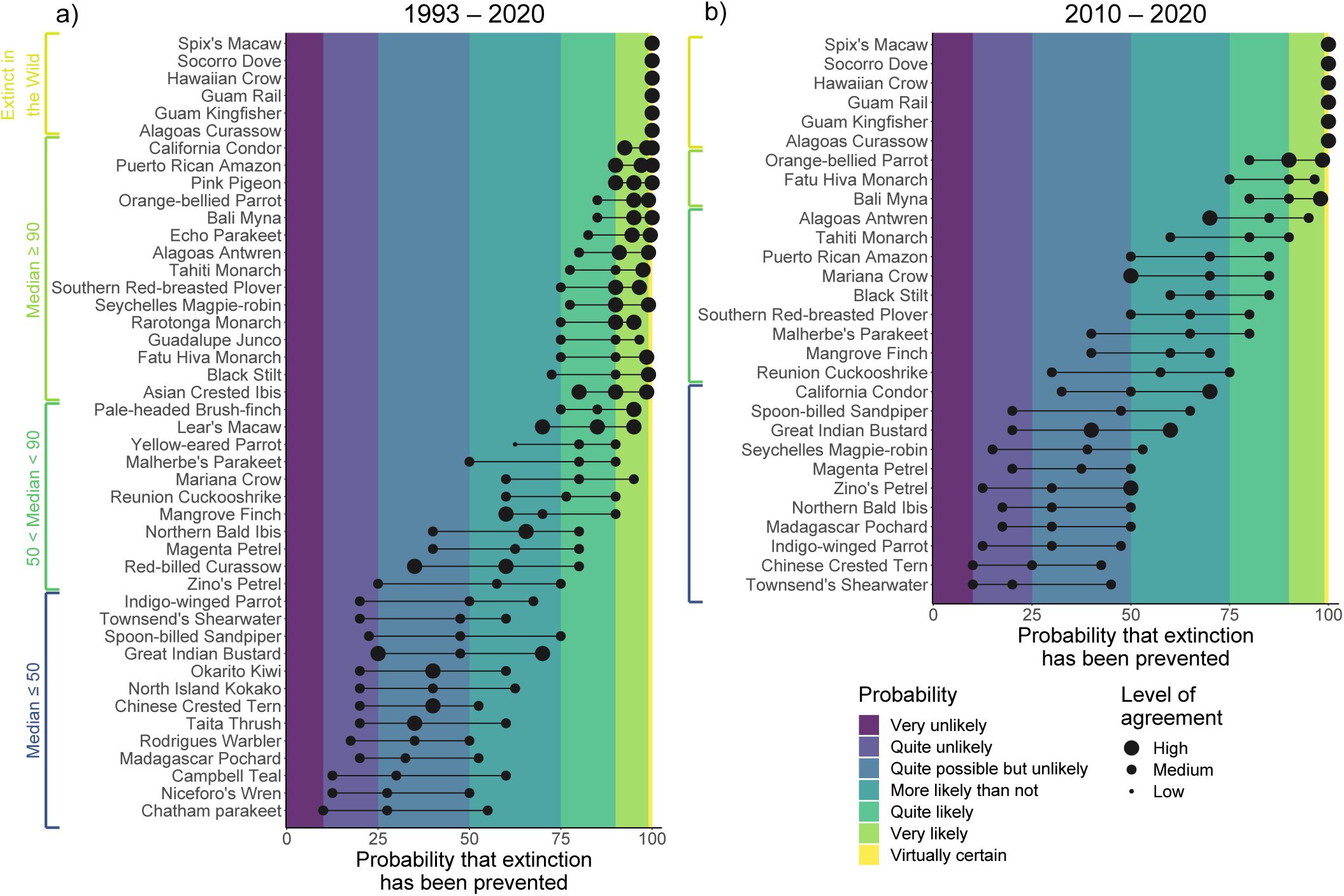
Probability that extinction of bird species would have occurred in the absence of conservation action during (a) 1993-2020 (N = 45 species) and (b) 2010-2020 (N = 29 species). Values represent medians calculated from estimates by 28 evaluators, except for species that are Extinct in the Wild, where it was set at 100%. For a description of the probability categories see Table S1, based on Keith et al. (2017). Guam Rail was assessed as Extinct in the Wild until 2016, but was reintroduced and assessed as Critically Endangered by 2019 (BirdLife International, 2020). We therefore set its probability to 100% for both time periods.

Of 23 candidate bird species for 2010-2020, three had a median best estimate ≥90% that their extinction was prevented (Fig. 1b), with a median best estimate >50% for a further nine species. Agreement among evaluators for these 12 species was high for one and medium for 11 species. Including six species listed as Extinct in the Wild, we consider that 9–18 bird species would have gone extinct without conservation during 2010-2020. In contrast, one bird species went extinct since 2010 (Table S2). Overall, the number of bird extinctions since 2010 would have been 10–19 times higher without conservation (10–19 vs. 1) (Table S3).

Of 21 candidate mammal species for 1993-2020, four had a median best estimate ≥90% that their extinction was prevented (Fig. 2a), and a further nine a median best estimate >50%. Agreement among evaluators for these 13 species was high for eight and medium for five species. Three species were listed as Extinct in the Wild during the time period (Table S4). Hence, we consider that 7–16 mammal species would have gone extinct without conservation during 1993-2020. Given that five mammal species are confirmed or suspected to have gone extinct since 1993 (Table S2), the number of mammal extinctions since 1993 would have been 2.4–4.2 times higher without conservation (12– 21 vs. 5) (Table S3).

**Figure 2.**
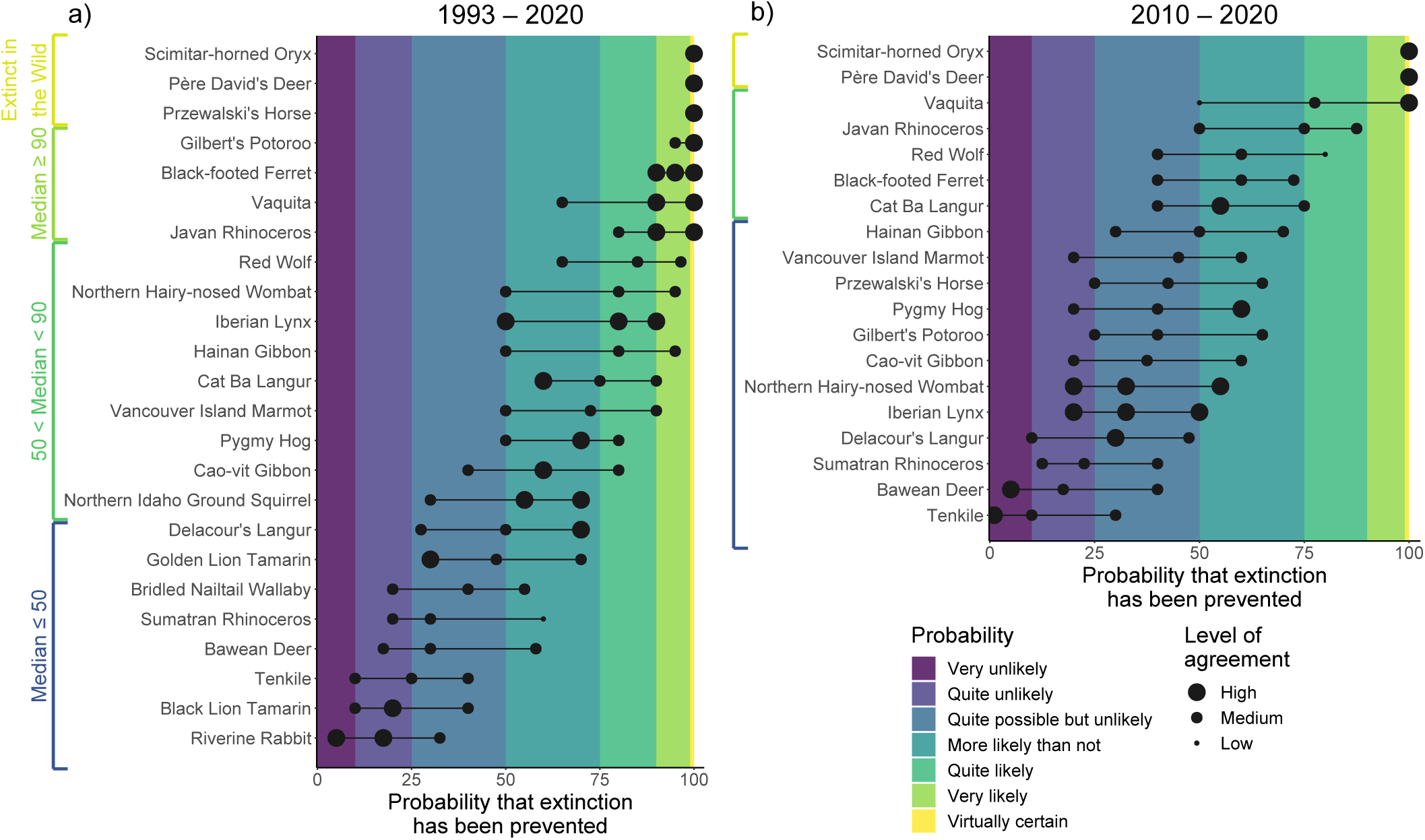
Probability that extinction of mammal species would have occurred in the absence of conservation action during (a) 1993-2020 (N = 24 species) and (b) 2010-2020 (N = 19 species). Values represent medians calculated from estimates by 26 evaluators, except for species that are Extinct in the Wild, where it was set at 100%. For a description of the probability categories see Table S1, based on Keith et al. (2017). Przewalski’s Horse was assessed as Extinct in the Wild in 1996, but was reintroduced and assessed as Critically Endangered by 2008. We therefore set its probability to 100% for 1993-2020, but asked evaluators to score for 2010-2020.

Of 17 candidate mammal species for 2010-2020, none had a median best estimate of ≥90% that their extinction was prevented and five had a median best estimate >50% (Fig. 2b). Agreement among evaluators for these five species was high for one and medium for four species. Including two species listed as Extinct in the Wild, we consider that 2–7 mammal species would have gone extinct without conservation during 2010-2020. No mammal species have been documented to have become extinct since 2010, so for this group all extinctions have been prevented by conservation.

These numbers of prevented extinctions are broadly consistent with values obtained following an approach of summing the median best estimates across all candidates (analogous to the approach for estimating the number of extinctions proposed by Akçakaya et al., 2017): 32.9 bird and 15.9 mammal species in 1993-2020, and 18.7 bird and 9.0 mammal species in 2010-2020.

The 32 identified bird species whose extinction was likely prevented during 1993-2020 occur (or occurred, for Extinct in the Wild species) in 25 countries, including six in New Zealand, five in Brazil, and three in Mexico (Fig. 3a); 65% are restricted to islands (excluding mainland Australia). The 16 identified mammal species occur in 23 countries, including five in China and three in Vietnam and the USA respectively (Fig. 3b); 19% are restricted to islands.

**Figure 3.**
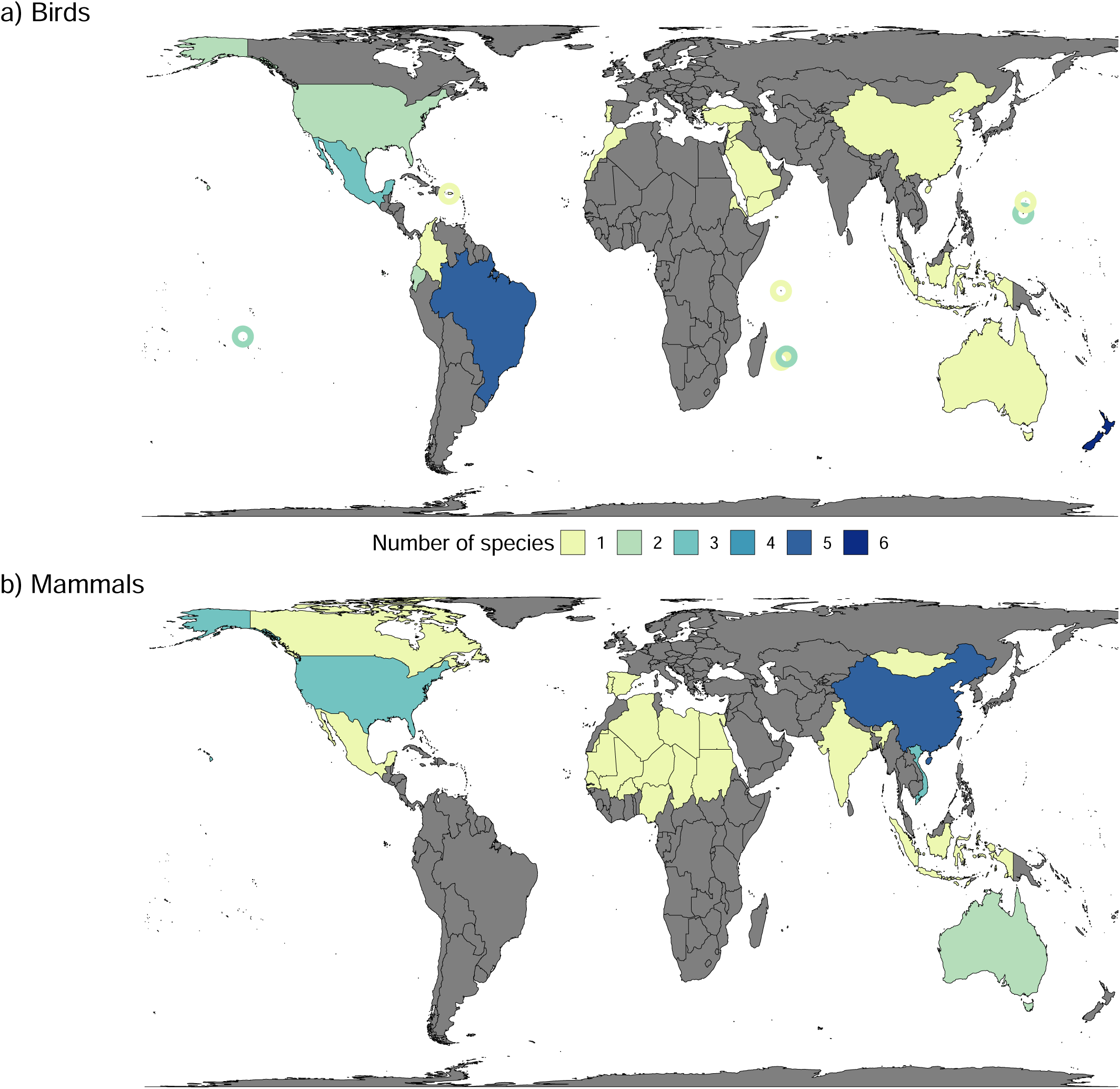
Number of (a) bird (N = 32) and (b) mammal (N = 16) species for which extinction is likely to have occurred (i.e. median probability >50%) in the absence of conservation action during 1993-2020, per country. Circles show small island nations and overseas territories, and are coloured according to the key. Species listed as Extinct in the Wild (IUCN, 2020) were mapped in the last countries where they occurred, or are presumed to have occurred.

Of the 32 identified bird species, 16% are currently classified as Extinct in the Wild, 47% as Critically Endangered, 28% as Endangered, and 9% as Vulnerable, with 53% having increasing or stable populations (Fig. 4a). Of the 16 identified mammal species, 13% are Extinct in the Wild, 56% Critically Endangered and 31% Endangered (Fig. 4b), with 31% having increasing or stable populations.

**Figure 4.**
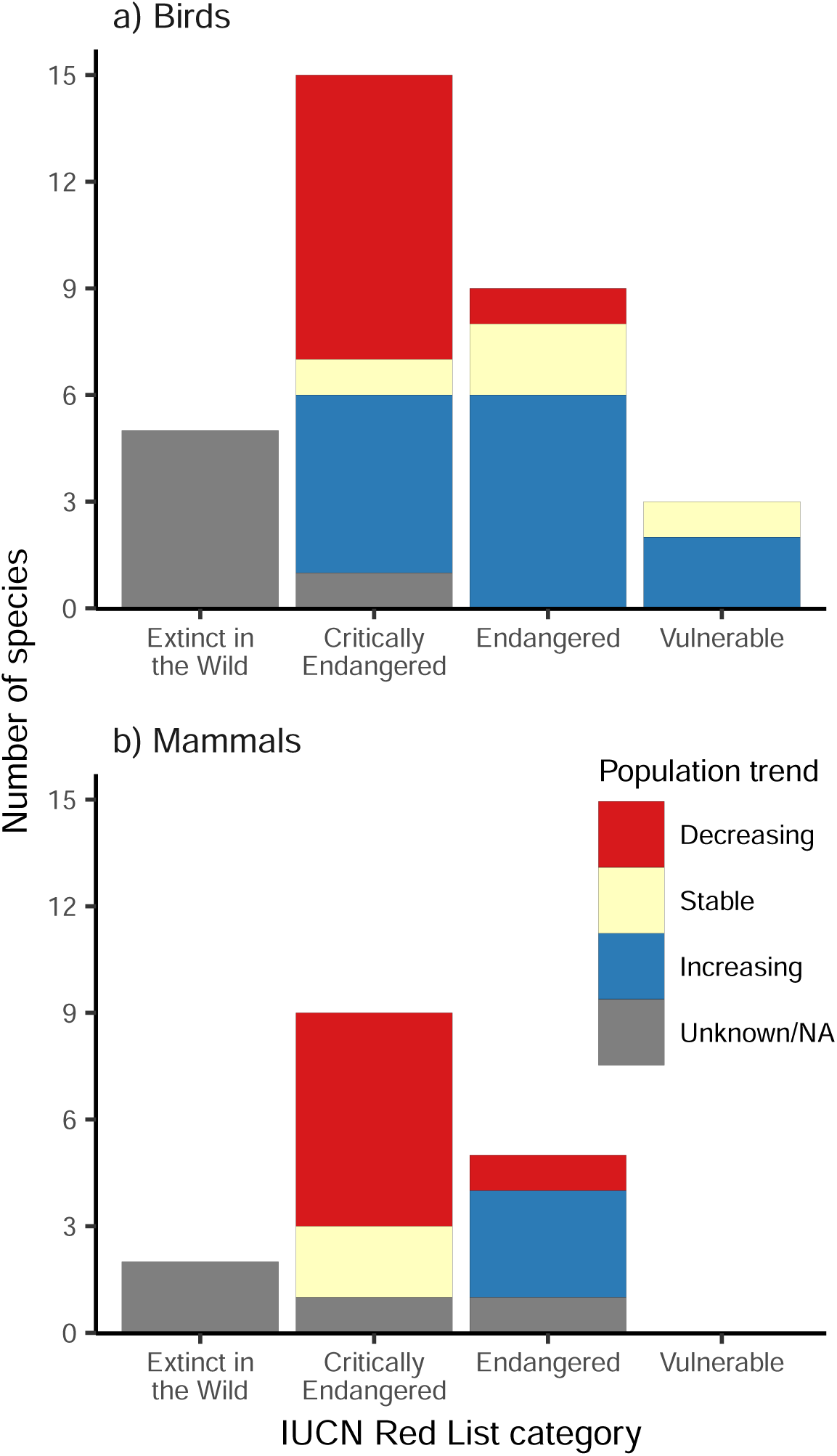
2019 IUCN Red List categories and population trends of (a) bird (N = 32) and (b) mammal (N = 16) species for which extinction is judged to have been likely (i.e. median probability >50%) to have occurred in the absence of conservation action, during 1993-2020.

The most frequent (including both current and past) threats to the 32 identified bird species are invasive species, followed by habitat loss through agriculture and aquaculture, and hunting (impacting 78%, 56% and 53% of species respectively) (Fig. 5a). The most frequent threats to the 16 identified mammal species are hunting, agriculture and aquaculture, and invasive species (impacting 75%, 75% and 50% of species respectively) (Fig. 5b).

**Figure 5.**
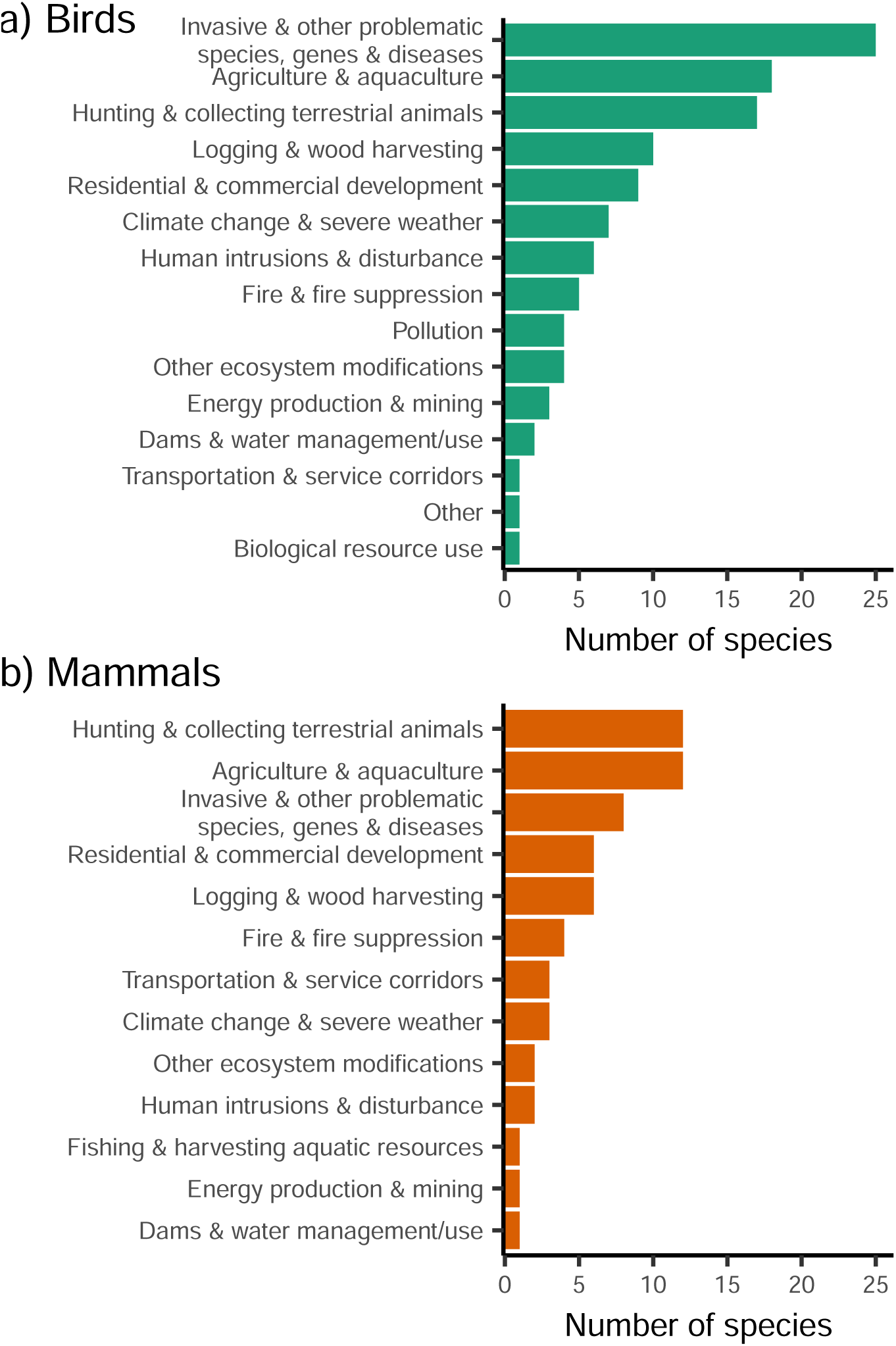
Current and past threats to (a) bird (N = 32) and (b) mammal (N = 16) species for which extinction is judged to have been likely (i.e. median probability >50%) to have occurred in the absence of conservation action during 1993-2020. Threats are taken from the IUCN threat classification scheme level 1 (Salafsky et al., 2008).

The most frequently implemented actions for the 32 identified bird species were invasive species control, ex-situ conservation, and site/area protection (for 66%, 63%, 59% of species respectively) (Fig. 6a). For the 16 mammal species, the most frequent actions were legislation, reintroductions, and ex-situ conservation (for 88%, 56%, 56% of species respectively) (Fig. 6b).

**Figure 6.**
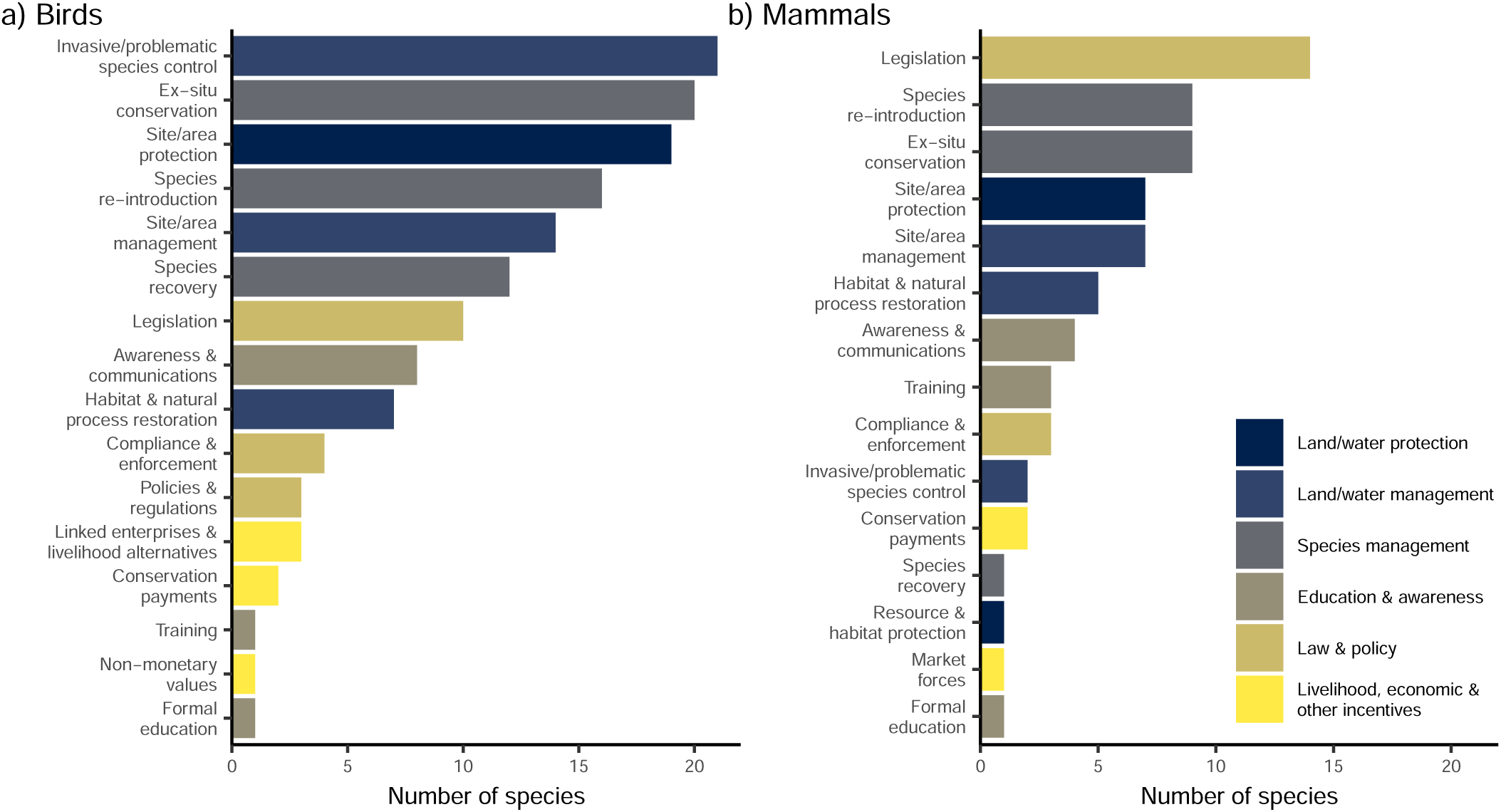
Conservation actions for (a) bird (N = 32) and (b) mammal (N = 16) species for which extinction is judged to have been likely (i.e. median probability >50%) to have occurred in the absence of conservation action during 1993-2020. Actions are taken from the IUCN action classification scheme level 2, while colours denote level 1 (Salafsky et al., 2008). Both in-situ and ex-situ actions are included for species that are Extinct in the Wild.

## Discussion

Our results indicate that the extinction of 28–48 bird and mammal species was prevented between 1993-2020, and of 11–25 bird and mammal species between 2010-2020. At the same time, there were 15 confirmed or strongly suspected bird and mammal extinctions since 1993, including one since 2010 (Alagoas Foliage-gleaner *Philydor novaesi*). Hence the number of extinctions would have been 2.9–4.2 times higher for 1993-2020, and 12–26 times higher for 2010–2020. Further extinctions since 2010 may come to light due to time-lags before detecting extinctions (Butchart et al., 2018). If the rate of extinctions observed in 1993-2009 (8.2/decade) is found to have continued during 2010-2020, the number of extinctions without conservation would still be two to four times higher (19.2-33.2 vs 8.2). Our counterfactual analyses therefore provide a strikingly positive message that conservation has substantially reduced extinction rates for birds and mammals.

Our analyses underestimate the impact of conservation in several ways. First, our process to identify candidate species may have failed to select all species whose extinction has been prevented. This could conceivably include some Endangered species that are rapidly declining (Criterion A) or with small, declining population sizes (Criterion C2). Others may have been missed owing to lack of information (for example, for Critically Endangered species tagged as Possibly Extinct, whose continued survival is uncertain). Second, we used the definition of extinction (the death of the last individual) adopted by IUCN (2012). Without conservation in the time periods we considered, a few additional species may well have become functionally extinct, such as the long-lived Okarito Kiwi *Apteryx rowi* (longevity ca. 100 years). Low numbers of old individuals of this species would have remained past 2020, even in the absence of ongoing measures to protect eggs and chicks from predation (BirdLife International 2017). Third, we considered only bird and mammal species, yet an additional 66 species in other taxa are listed as Extinct in the Wild on the IUCN Red List (IUCN 2020). These would be extinct without ex-situ efforts, while other extant species would have gone extinct without in-situ efforts. Lastly, we examined only species at the brink of extinction: many more species would have deteriorated in conservation status in the absence of conservation (Hoffmann et al., 2015). We also note the influence of stochastic processes on our results: Puerto Rican Amazon *Amazona vittata* scored highly because of the occurrence of a hurricane that left only a population of birds in the wild derived from captive breeding, while Madagascar Pochard *Aythya innotata*, with an even smaller wild population, scored low because no such stochastic event occurred.

Conversely, not all the species we identified as prevented extinctions can be described as conservation successes. For some, conservation has slowed declines but perhaps not sufficiently to prevent near-future extinction. This may be the case for the Vaquita *Phocoena sinus*, of which six individuals were known to remain in September 2018 (Jaramillo-Legorreta et al., 2019). Conservation may have slowed the catastrophic decline but is proving unable to halt it.

The conservation actions we found to have contributed to avoiding species extinctions echo the respective main threats. The most frequent threat for birds was invasive species, and management of invasive species was the key response. For mammals, the prominence of legislation as a conservation action likely reflects efforts to curb the main threat of hunting and collecting. Site/area protection are featured frequently as actions for both taxa, considering that agriculture and aquaculture, logging, and residential development appear as persistent threats. The importance of ex-situ conservation and species reintroductions reflects the large numbers of species whose persistence has relied on captive-bred populations, sometimes completely (for the Extinct in the Wild species, Table S4), others as a source of individuals for translocations and population reinforcements (Table S6). Two formerly Extinct in the Wild species have been reintroduced successfully since 1993: Przewalski’s Horse *Equus ferus* and Guam Rail *Hypotaenidia owstoni*.

Assessing the probability that species would have gone extinct under a counterfactual scenario inherently involves a degree of uncertainty. Judgements are more certain when better information is available on the species, and it is possible that we missed some information that would have changed the probabilities assigned to species. We attempted to minimise this risk by starting from the comprehensive IUCN Red List assessments, incorporating up to date information from 124 species experts, and asking each evaluator to examine more thoroughly a small subset of species prior to the calls. We undertook two calls per taxon, with largely different sets of evaluators per call. As slightly different information was discussed during each call, there were some differences in probability estimates between calls. To reduce this effect, we relayed information gained during the first call to evaluators on the second call, but in some cases new information came to light during the second call (see Supplementary Information). However, differences between calls had little effect on the overall results. Two mammal species had an overall probability ≤50%, but would be included (i.e. an estimate >50%) based on scores from the second call only, and two bird and one mammal species had an overall probability >50%, but would be excluded (i.e. an estimate ≤50%) based on scores from the second call only.

Our results show that despite the ongoing loss of biodiversity, a substantial number of extinctions was prevented since the inception of the CBD. While Aichi Target 12 has not been met, the rate of extinctions since it was adopted would have been at least twice as high (and potentially an order of magnitude higher) without conservation action. These actions were implemented by a combination of governments, NGOs, zoos, scientists, volunteers and others. Nevertheless, the species we identified remain highly threatened, and most require continued substantial conservation investment to ensure their survival. Given the ongoing scale and projected growth in pressures on biodiversity (IPBES 2019), considerably greater efforts are needed to prevent the extinction and improve the status of the 6,413 species currently assessed as Critically Endangered on the IUCN Red List (IUCN 2020). Our results should motivate the world’s governments currently negotiating goals and targets on nature conservation in the CBD’s post-2020 biodiversity framework to redouble their commitments to prevent extinctions. Not only is this hugely important (Gascon et al., 2015) but also, as we have demonstrated here, eminently feasible.

## Acknowledgements and data

We thank the following species experts for providing input to this project: Antonio Ortiz Alcaraz, Giovanni Amori, Arthur Barbosa Andrade, Helder Farias Pereira de Araujo, Andrew Bamford, António Eduardo Araújo Barbosa, Gobind Sagar Bhardwaj, Caroline Blanvillain, Luca Borghesio, Chris Bowden, Lee Boyd, Dave Boyle, Amedee Brickey, Rachel M. Bristol, Andrew Burbidge, Ian Burfield, Rick Camp, Fernando Solis Carlos, Kevin Carter, Simba Chan, Susan Cheyne, Francesca Cunninghame, Dave Currie, Pedro Ferreira Develey, Andrew Digby, John Dowding, Márcio Amorim Efe, Jorge Fernández Orueta, Julie Gane, Thomas Ghestemme, Mwangi Githiru, Amanda Goldberg, Andrew Grant, Rhys Green, Terry Greene, Rod Hitchmough, Alan Horsup, Simon Hoyle, John Hughes, John Innes, Todd Katzner, Jonathan Kearvell, Bruce Kendall, Cecilia Kierulff, Andrew Legault, Neahga Leonard, Jorgelina Marino, Juan Esteban Martínez, Pete McClelland, Neil McCulloch, Michael McMillan, Patricia Moehlman, Julio Hernández Montoya, Tilo Nadler, Steffen Oppel, Antonio Ortiz, Oliver Overdyck, Erica C. Pacifico, Phil Palmer, Fernando C. Passos, Erica Perez, Benjamin Timothy Phalan, Mike Phillips, Huy Hoàng Qu⍰c, Lily-Arison Réné De Roland, Johanna Rode-Margono, Carlos Ramon Ruiz, Michael J. Samways, H. Martin Schaefer, Jessica Scrimgeour, Gono Semiadi, Claudio Sillero-Zubiri, Herminio Alfredo Leite Silva Vilela, Luis Fábio Silveira, Elenise Angelotti Bastos Sipinski, Fernando Solis, Christine Steiner São Bernardo, Bibhab Talukdar, Vikash Tatayah, Bernie Tershy, Cobus Theron, Jean-Claude Thibault, Jeff J. Thompson, Sam Turvey,, Thomas White, Peter Widmann, Yu Xiao-Ping, Ding Li Yong, Glyn Young, Francis Zino, Christoph Zöckler. ARP is supported by FAPESP (Fundação de Pesquisa do Estado de São Paulo) and CNPq (Conselho Nacional de Desenvolvimento Científico e Tecnológico). All code and data can be found at http://github.com/rbolam/Prevented_bird_and_mammal_extinctions. This research has been granted approval by the Newcastle University Ethics Committee (Reference 15388/2018).

## References

Akçakaya, H.R., Keith, D.A., Burgman, M., Butchart, S.H., Hoffmann, M., Regan, Boakes E. (2017). Inferring extinctions III: A cost-benefit framework for listing extinct species. Biological Conservation, 214, 336–342. doi: 10.1016/j.biocon.2017.07.027

Akçakaya, H.R., Bennett, E.L., Brooks, T.M., Grace, M.K., Heath, A., Hedges, S., Mallon, D.P. (2018). Quantifying species recovery and conservation success to develop an IUCN Green List of Species. Conservation Biology, 32(5), 1128–1138. doi: 10.1111/cobi.13112

BirdLife International (2017). Apteryx rowi. The IUCN Red List of Threatened Species 2017: e.T22732871A119169794. Retrieved 05/02/2020 from https://dx.doi.org/10.2305/IUCN.UK.2017-3.RLTS.T22732871A119169794.en

BirdLife International (2019). Hypotaenidia owstoni. The IUCN Red List of Threatened Species 2019: e.T22692441A156506469. Retrieved 10/02/2020 from https://dx.doi.org/10.2305/IUCN.UK.2019-3.RLTS.T22692441A156506469.en

Butchart, S.H., Stattersfield, A.J. and Collar, N.J. (2006). How many bird extinctions have we prevented? Oryx, 40(3), 266-278. doi: 10.1017/S0030605306000950

Butchart, S. H. M., Akçakaya, H. R., Chanson, J., Baillie, J. E. M., Collen, B., Quader, S., Mace, G. M. (2007) Improvements to the Red List Index. PLoS ONE 2: e140. doi: 10.1371/journal.pone.0000140

Butchart, S.H., Lowe, S., Martin, R.W., Symes, A., Westrip, J.R. and Wheatley, H. (2018). Which bird species have gone extinct? A novel quantitative classification approach. Biological Conservation, 227, 9–18. doi: https://doi.org/10.1016/j.biocon.2018.08.014

CBD (2014). Global Biodiversity Outlook 4. Montréal, 155 pages.

Diamond, I.R., Grant, R.C., Feldman, B.M., Pencharz, P.B., Ling, S.C., Moore, A.M. & Wales, P.W. (2014). Defining consensus: a systematic review recommends methodologic criteria for reporting of Delphi studies. Journal of Clinical Epidemiology, 67, 401–409. doi: 10.1016/j.jclinepi.2013.12.002

Díaz, S., Settele, J., Brondízio, E.S., Ngo, H.T., Agard, J., Arneth, A., Garibaldi, L.A. (2019). Pervasive human-driven decline of life on Earth points to the need for transformative change. Science, 366(6471). doi: 10.1126/science.aax3100

Gascon, C., Brooks, T.M., Contreras-MacBeath, T., Heard, N., Konstant, W., Lamoreux, J., Al Mubarak, R.K. (2015). The importance and benefits of species. Current Biology, 25(10), R431–R438. doi: 10.1016/j.cub.2015.03.041

Hemming, V., Burgman, M.A., Hanea, A.M., McBride, M.F. and Wintle, B.C. (2018). A practical guide to structured expert elicitation using the IDEA protocol. Methods in Ecology and Evolution, 9(1), 169-180. doi: 10.1111/2041-210X.12857

Hoffmann, M., Hilton-Taylor, C., Angulo, A., Böhm, M., Brooks, T. M., Butchart, S. H. M., … Stuart, S. N. (2010). The impact of conservation on the status of the world’s vertebrates. Science 330, 1503–1509. doi: 10.1126/science.1194442

Hoffmann, M., Belant, J.L., Chanson, J.S., Cox, N.A., Lamoreux, J., Rodrigues, A.S., … Stuart, S.N. (2011). The changing fates of the world’s mammals. Philosophical Transactions of the Royal Society B: Biological Sciences, 366(1578), 2598–2610. doi: 10.1098/rstb.2011.0116

Hoffmann, M., Duckworth, J.W., Holmes, K., Mallon, D.P., Rodrigues, A.S. and Stuart, S.N. (2015). The difference conservation makes to extinction risk of the world’s ungulates. Conservation Biology, 29(5), 1303–1313. doi: 10.1111/cobi.12519

IPBES (2019): Summary for policymakers of the global assessment report on biodiversity and ecosystem services of the Intergovernmental Science-Policy Platform on Biodiversity and Ecosystem Services. S. Díaz, J. Settele, E. S. Brondízio E.S., H. T. Ngo, M. Guèze, J. Agard, A. Arneth, P. Balvanera, K. A. Brauman, S. H. M. Butchart, K. M. A. Chan, L. A. Garibaldi, K. Ichii, J. Liu, S. M. Subramanian, G. F. Midgley, P. Miloslavich, Z. Molnár, D. Obura, A. Pfaff, S. Polasky, A. Purvis, J. Razzaque, B. Reyers, R. Roy Chowdhury, Y. J. Shin, I. J. Visseren-Hamakers, K. J. Willis, and C. N. Zayas (eds.). IPBES secretariat, Bonn, Germany. 56 pages.

IUCN (2019). The IUCN Red List of Threatened Species. Version 2019-1. Retrieved 24/01/2019 from https://www.iucnredlist.org

IUCN (2020). The IUCN Red List of Threatened Species. Version 2019-3. Retrieved 05/02/2020 from https://www.iucnredlist.org

Jaramillo-Legorreta, A.M., Cardenas-Hinojosa, G., Nieto-Garcia, E., Rojas-Bracho, L., Thomas, L., Ver Hoef, J.M., … Tregenza, N. (2019). Decline towards extinction of Mexico’s vaquita porpoise (Phocoena sinus). Royal Society open science, 6(7), 190598. doi: 10.1098/rsos.190598

Keith, D.A., Butchart, S.H., Regan, H.M., Harrison, I., Akçakaya, H.R., Solow, A.R. and Burgman, M.A. (2017). Inferring extinctions I: A structured method using information on threats. Biological Conservation, 214, 320–327. doi: 10.1016/j.biocon.2017.07.026

Knol, A.B., Slottje, P., van der Sluijs, J.P. and Lebret, E. (2010). The use of expert elicitation in environmental health impact assessment: a seven step procedure. Environmental Health, 9(1), 19. doi: 10.1186/1476-069X-9-19

Martin, T.G., Burgman, M.A., Fidler, F., Kuhnert, P.M., Low-Choy, S., McBride, M. and Mengersen, K. (2012). Eliciting expert knowledge in conservation science. Conservation Biology, 26(1), 29–38. doi: 10.1111/j.1523-1739.2011.01806.x

Morgan, M.G. (2014). Use (and abuse) of expert elicitation in support of decision making for public policy. Proceedings of the National Academy of Sciences, 111(20), 7176–7184. doi: 10.1073/pnas.1319946111

Mukherjee, N., Huge, J., Sutherland, W.J., McNeill, J., Van Opstal, M., Dahdouh-Guebas, F. and Koedam, N. (2015). The Delphi technique in ecology and biological conservation: applications and guidelines. Methods in Ecology and Evolution, 6(9), 1097–1109. doi: 10.1111/2041-210X.12387

Pacifici, M., Santini, L., Di Marco, M., Baisero, D., Francucci, L., Marasini, G.G., … Rondinini, C. (2013). Generation length for mammals. Nature Conservation, 5, 89. doi: 10.3897/natureconservation.5.5734

Salafsky, N., Salzer, D., Stattersfield, A.J., Hilton-Taylor, C., Neugarten, R., Butchart, S.H.M., … Wilkie, D. (2008). A standard lexicon for biodiversity conservation: unified classifications of threats and actions. Conservation Biology, 22(4), 897–911. doi: 10.1111/j.1523-1739.2008.00937.x

Szabo, J.K., Butchart, S.H., Possingham, H.P. and Garnett, S.T. (2012). Adapting global biodiversity indicators to the national scale: A Red List Index for Australian birds. Biological Conservation, 148(1), 61–68. doi: 10.1016/j.biocon.2012.01.062

Timmins, R.J., Hedges, S. & Robichaud, W. (2016). Pseudoryx nghetinhensis. The IUCN Red List of Threatened Species 2016: e.T18597A46364962. Retrieved 30/01/2020 from https://dx.doi.org/10.2305/IUCN.UK.2016-2.RLTS.T18597A46364962.en

Troudet, J., Grandcolas, P., Blin, A., Vignes-Lebbe, R. and Legendre, F. (2017). Taxonomic bias in biodiversity data and societal preferences. Scientific reports, 7(1). doi: 10.1038/s41598-017-09084-6

von der Gracht, H.A. (2012). Consensus measurement in Delphi studies. Review and implications for future quality assurance. Technological Forecasting and Social Change, 79, 1525–1536. doi: 10.1016/j.techfore.2012.04.013

Walsh, J.C., Watson, J.E., Bottrill, M.C., Joseph, L.N. and Possingham, H.P. (2013). Trends and biases in the listing and recovery planning for threatened species: an Australian case study. Oryx, 47(1), 134–143. doi: 10.1017/S003060531100161X

Young, R.P., Hudson, M.A., Terry, A.M.R., Jones, C.G., Lewis, R.E., Tatayah, V., Butchart, S.H.M. (2014). Accounting for conservation: Using the IUCN Red List Index to evaluate the impact of a conservation organization. Biological Conservation, 180, 84–96. doi: 10.1016/j.biocon.2014.09.039

