## Supplementary material for "How many bird and mammal extinctions has recent conservation action prevented?"

### Online supporting information

#### Extended methods

##### Initial screening process

We examined which bird and mammal species may have been prevented from going extinct during two time periods: 1993–2020, and 2010–2020. We chose these years as they correspond, respectively: to the period in which the Convention on Biological Diversity (CBD) has been in force; and the period for implementation of the CBD Strategic Plan for Biodiversity with its 20 Aichi targets (starting in 2011).

We downloaded the Red List assessment history of birds and mammals from the IUCN Red List on 24 January 2019 (IUCN 2019). We considered species as potential candidates if they are currently extant or Extinct in the Wild and were, at any point since 1993:

- listed as Extinct in the Wild
- listed as Critically Endangered (any criterion)
- listed as Endangered under criterion D, i.e., with fewer than 250 mature individuals

In some cases, species' Red List assessments in any given year are subsequently found to have been erroneous, either because of new information (e.g., a larger or smaller population size than previously estimated) or because of taxonomic changes (e.g., a species split in two). In these cases, the categories used in the above-mentioned filtering process were the ones obtained after retrospective correction based on current knowledge rather than the initial Red List classification (as also applied in calculating the IUCN Red List Index; Butchart et al 2007, Hoffmann et al., 2011). This initial list included 368 bird and 263 mammal species. Of these, six bird and two mammal species were classified as Extinct in the Wild as of 2019 (IUCN 2019) and considered to have a 100% probability that they would have gone extinct without further conservation.

We then narrowed down the remaining list of species by examining the IUCN Red List accounts of these species. We retained species for which there was evidence that it could have gone extinct during the time periods under consideration: those with < 200 individuals or those with populations with very rapid declines, provided there was a significant threat or suite of threats that might have driven them extinct in the absence of actions, and for which actions were implemented that mitigated these threats (Butchart et al., 2006). Hence, we excluded species with no information on conservation actions, or for which actions appeared insufficient to address the main threats. Where the effectiveness of actions was unclear based on the information available, the species was retained.

To give some examples, Saola *Pseudoryx nghetinhensis* is severely threatened by poaching, but the Red List account details that protected areas and anti-poaching measures are largely ineffective (Timmins et al., 2016), hence it was excluded. This decision is supported by the conclusion of Hoffmann et al. (2015) who considered an Extinct listing for this species as a pessimistic scenario, noting that "while Saola has been found in several protected areas in both its range countries, much potential habitat is outside protected areas. Moreover the PAs are not sufficiently well managed to remove major hunting threats to this species or to have headed off major habitat conversion that would otherwise have occurred. Therefore, this species would, following cessation of conservation actions in 1996, have continued its decline towards extinction in a similar manner to that seen in the last 12 years". Some notable conservation success stories such as Black Robin *Petroica traversi* or

Mauritius Kestrel *Falco punctatus* were also excluded, as actions took place for these species prior to 1993 and they had recovered sufficiently by 1993 to not be considered by us.

This filtering process was undertaken separately by two groups of people: all species were screened by a team at Newcastle University (either by FB, LM or PJKM). Birds were additionally screened by CH, RM, and HW at BirdLife International, and mammals were additionally screened by MA at the Global Mammal Assessment. The lists were then compared. Initial agreement for categories was 71% for birds and 80% for mammals. The Newcastle team then investigated the species with different assessments in more detail, and either confirmed the assessment by the other teams (based on the new information they provided) or added more information to support their assessment. This information then went back to the teams at BirdLife International or the Global Mammal Assessment, respectively, and any remaining differences were discussed. Species for which consensus could not be achieved were retained on the candidate list. The shortlist of species included 48 bird and 25 mammal species (in addition to the six Extinct in the Wild bird and two Extinct in the Wild mammal species).

#### Compiling species information

Data were extracted for all initial candidate species. The IUCN Red List species accounts were searched to extract population estimates and trends for 1993, 2010, and the latest assessment year, as well as threats and conservation actions. Information on threats and actions were extracted for the respective time periods. We also added information on generation lengths of birds based on BirdLife International (2020), and generation lengths and longevity for mammals, based on Pacifici et al. (2013). We added a summary of the reasons why each species was included as a candidate for each period.

The initial 48 bird and 25 mammal accounts were reviewed by experts on the species, as identified by BirdLife International for birds, and by the Global Mammal Assessment for mammals. We contacted 197 bird and 77 mammal experts. For birds, we received 88 responses for 45 species (94%). For mammals, we received 36 responses for 24 species (96%). In some cases, the new information provided by experts led us to then exclude the species, narrowing our list further to 39 bird and 21 mammal candidate species. Whereas all of these were considered as candidates for potentially having gone extinct without conservation action during 1993-2020, a shorter list of 23 bird and 17 mammal species was considered for the 2010-2020 period. This was typically because the population had become sufficiently large, widely distributed, and/or secure by 2010 that it was implausible that the species would have gone extinct if action had ceased in 2010). One exception to this was Przewalski's Horse *Equus ferus.* This species was Extinct in the Wild in 1993, and was included with other species that were assessed as Extinct in the Wild for 1993 - 2020. However, it was assessed as Critically Endangered in 2008 following successful reintroduction and we therefore included it as a candidate for the 2010-2020 time period.

#### Delphi exercise

The next step involved assigning probabilities that conservation action prevented extinction in the candidate species, which was done by species conservation experts following a Delphi protocol (accessible at https://osf.io/rk4ep/).

##### *Selection of evaluators*

Potential evaluators were identified from the contributors to Red List Assessments for birds by the Red List Authority for birds (BirdLife International) and for mammals by the Global Mammal Assessment team (who coordinate Red List assessments for mammals) based on the criteria set out below:

i) Experience of species monitoring and conservation, and an understanding of the challenges of searching for and detecting rare species AND
ii) This experience spans multiple species AND
iii) Experience of quantifying estimates based on uncertain information (e.g. through undertaking Red List assessments)

Those identified included individuals from a diversity of backgrounds, nationalities, regions, and gender. A total of 66 evaluators meeting these criteria were invited to participate (31% female, and 30% based in the Global South), of whom 11 were invited to participate in both exercises. 28 bird and 26 mammal evaluators took part, of which 34% were female, and 23% based in the Global South. The following authors took part in the Delphi exercise for birds: FCB, LM, CH, MH, RWM, PJKM, ASLR, HW, YBG, MFC, PAC, BF, SG, JJG, JFL, ACL, LL, SPM, DPM, FMS, LMR, MCR, RJS, PS, TS, JRSW, RPY, SHMB. The following authors took part in the Delphi exercise for mammals: FCB, LM, TMB, MH, PJKM, ASLR, CR, JC, MFC, CRD, DOF, CNJ, RJK, SRBK, JFL, DPM, EM, ARP, TJR, NSR, LR, EVD, PV, JCZW, RPY, SHMB.

##### *Preparation and conducting the Delphi exercise*

Ethical approval to undertake this activity was given by Newcastle University, reference number 15388/2018. We followed the IDEA protocol (Investigate, Discuss, Estimate and Aggregate) in which experts each make an independent, anonymous, quantitative assessment, followed by facilitated discussion, followed by another quantitative assessment, followed by aggregation of their estimates (Hemming et al., 2018). In order to do this, each species was evaluated by all bird or all mammal evaluators. A few evaluators are more involved with the direct management of some species under consideration, and they might be incentivised to assign high probabilities to those species. However, as their score was only one out of 28 for bird species, or one out of 26 for mammal species, we did not consider this to be an issue.

Evaluators received instructions and background information for the Delphi technique (see *Information circulated to evaluators* at the end of this document). The evaluators were sent the same list of candidate species for birds or mammals, depending on whether they were identified as evaluators for birds, mammals, or both groups. Each evaluator received lists in which the species appeared in a different, random order. For each species, the evaluators received the compiled and revised species information.

The questions were based on Morgan (2014) and Hemming et al. (2018), with questions on extreme values asked first to avoid anchoring (Morgan, 2014). When using qualitative terms to describe probabilities, there are large differences between individuals in the perceived probability (Morgan, 2014), so we used numerical values instead. To ensure that our questions and background information on the exercise were not ambiguous, we tested them on eight non-species expert students/staff at the School of Natural and Environmental Sciences at Newcastle University, UK, based on five sample species and using the background information prepared for the evaluators. We took note of any arising questions. The students and staff were asked what their understanding of the questions was, to ensure our intended meaning was clear to everyone. The information for evaluators was revised after the exercise. The three questions for all species were:

Realistically, what do you think is the **(1) lowest plausible probability/ (2) highest plausible probability/ (3) best estimate for the probability** that conservation action prevented extinction for this species during the period (i.e. what is the probability that, **if action had ceased in 1993**, and no subsequent actions were implemented, the species would have gone extinct by 2020)?

Additionally, if species met the criteria to be included for the time period 2010 – 2020, the three following questions were also asked:

Realistically, what do you think is the **the (4) lowest plausible probability/ (5) highest plausible probability/ (6) best estimate for the probability** that conservation action prevented extinction for this species during the period (i.e. what is the lowest plausible probability that, **if action had ceased in 2010**, and no subsequent actions were implemented, the species would have gone extinct by 2020)?

We explained that 'conservation action' encompassed the full range of interventions, including protected area establishment and management, legislation (e.g. to prohibit hunting), control or management of invasive alien species, control of hunting/trapping, habitat restoration, and species recovery interventions such as captive breeding, translocation, supplementary feeding, nest-site provision etc. We further clarified that the probabilities should be given by considering if all actions had ceased, including the degazettement of protected areas, and discontinuing captive breeding programmes (we include private collections here too). We recognise in practice, it might not be likely that all actions would cease, for example because there are legal implications in degazetting protected areas.

We also explained that we wanted the scores to reflect whether the species would have gone extinct in the wild if not for conservation action, meaning that the last individual in the wild would have disappeared by the beginning of 2020. Species that are listed as Extinct in the Wild on the IUCN Red List are listed separately in Tables S2 and S3.

We used a number of measures to reduce the attrition rate of evaluators: we used an Excel spreadsheet for the scoring to make this easy for everyone and made the species information available online. We piloted the exercise beforehand with a team of people to ensure the questions were not ambiguous and to estimate the time it would take for evaluators to make their scores. We also selected evaluators who have relevant expertise and interest as defined by our criteria above, and minimised the time between the iterations of the Delphi process, which was no more than six weeks between sending out instructions initially and revising the scores (Mukherjee et al., 2015).

##### *Measuring consensus*

Based on the results first returned by evaluators, we calculated the median lowest (question 1), highest (question 2) and best estimate (question 3) of probabilities that extinction has been prevented for each species (von der Gracht, 2012), for both time periods where applicable. We used medians, as unweighted approaches to combining expert knowledge are usually as accurate as more complex, weighted approaches (Knol et al., 2010; Martin et al., 2012), and chose medians to avoid undue influence from any outliers. To measure agreement, we defined seven classes of probability, based on Keith et al. (2017): very unlikely, quite unlikely, quite possible but unlikely, more likely than not, quite likely, very likely, virtually certain (see Table S1). We considered there to be high agreement if >50% of evaluators had placed their estimates within the same class, medium agreement if >50% of evaluators had placed their estimates within two adjacent classes, and low agreement for all other cases.

*Table S1. Range of probabilities and their meaning for whether extinction was prevented through conservation action (adapted from Keith et al., 2017).*

| **Range of probabilities** | *Was extinction prevented through conservation actions?* |
| --- | --- |

| | |
|---|---|
| 0.99 - 1.00 | The actions are **virtually certain** to have prevented extinction, i.e. would have prevented extinction in 99 of 100 species similar to the target. There is a less than a one in a hundred chance that the taxon would have persisted without conservation action during the period. |
| 0.90 - 0.98 | The actions are **very likely** to have prevented extinction, i.e. would have prevented extinction in 49 of 50 to 19 of 20 similar species. There is a one in fifty to one in twenty chance that the taxon would have persisted without conservation action during the period. |
| 0.75 - 0.89 | The actions are **quite likely** to have prevented extinction, i.e. would have prevented extinction in 19 of 20 to three in four similar species. There is a one in twenty to one in four chance that the taxon would have persisted without conservation action during the period. |
| 0.50 - 0.74 | The actions are **more likely than not** to have prevented extinction, i.e. would have prevented extinction of three-quarters of similar species. There is a one in four to 50:50 chance that the taxon would have persisted without conservation action during the period. |
| 0.25 - 0.49 | The actions are **quite possible but unlikely** to have prevented extinction, i.e. would have prevented extinction in one quarter to one half of similar species. There is more than a 50:50 and up to a 3 in 4 chance that the taxon would have persisted without conservation action during the period. |
| 0.10 - 0.24 | The actions are **quite unlikely** to have prevented extinction, i.e. would have prevented extinction of one tenth to one quarter of similar species. There is a 3 in 4 to 9 in 10 chance that the taxon would have persisted without conservation action during the period. |
| 0 - 0.09 | The actions are **very unlikely** to have prevented extinction, i.e. would have prevented extinction of up to one tenth of similar species. There is more than a 9 in 10 chance that the taxon would have persisted without conservation action during the period. |

##### *Scoring and discussion*

All evaluators scored all species independently first. We then shared with all evaluators the median and agreement of scores for each species, for each period. This was followed by teleconference video calls where we discussed each species in turn. We organised two calls for each of birds and mammals (to keep the group size sufficiently small to ensure all could contribute, and to address the evaluators' different time zones). Prior to the calls, each species was randomly assigned to two evaluators who were asked to familiarise themselves in more depth with the documentation provided and encouraged to seek any additional information online or offline and consider potential counterfactuals. During the first call, evaluators worked though the list in alphabetical order, and in reverse alphabetical order for the second call.

During the calls, each species was discussed in turn for no more than 10 minutes, with the discussion facilitated by one chair (SHMB for birds; MH for mammals). Facilitation included prompting to consider counterfactuals, choosing contrasting results for discussion and exploring potential reasons for contrasting results (Hemming et al., 2018). The median and degree of agreement of scores for each of the two periods were considered in the discussion (but individual scores remained anonymous). In a few cases where key information was mentioned during the first call which had not been part of the documentation on the species, this information was shared by the chair during the second call.

Following discussion of each species, all evaluators independently and confidentially re-estimated the probability that extinction had been prevented for each of the two time periods.

##### Subsequent analysis

The revised scores from the calls were used to recalculate median probabilities and agreement between evaluators (as defined above) for each time period, and these were used in the subsequent analysis. We summarised the overall results in terms of the number of species whose extinction has been prevented as X-Y, with X representing species with a median best estimate ≥90% (i.e. very likely to have had their extinction prevented) and Y representing species with a median best estimate > 50% (i.e. more likely than not to have had their extinction prevented), following an analogous approach for defining Extinct and Critically Endangered (Possibly Extinct) species (Butchart et al., 2018). We compared these total numbers with the rate at which extinctions have been prevented. We also calculated the latter rate as the sum of all best median probabilities across all candidate species (without setting thresholds), following an analogous approach for estimating the rate of extinction suggested by Akçakaya et al. (2018).

We plotted median probabilities for all candidate species for both time periods, for the lowest, best and highest estimate of the probability. We compared the scoring between calls for the best estimate for each species under each relevant time period using Kruskal-Wallis tests, as not all were normally distributed.

We mapped the current native or reintroduced distribution of the species for each country, except for Extinct in the Wild species, for which we mapped those countries in which they were native prior to extinction in the wild. We plotted threats, conservation actions, current Red List category and population trend, including for Extinct in the Wild species. We plotted threats according to IUCN threat level 1 (Salafsky et al., 2008), except for the threats Biological Resource Use and Natural Systems Modifications, as they comprise distinct threats. Biological Resource Use was therefore split into the relevant level 2 threats, namely Hunting and collecting terrestrial animals, Logging & wood harvesting, and Fishing & harvesting aquatic resources. The category Natural System Modifications was split into Fire and fire suppression, Dams and water management/use, and Other ecosystem modifications. We included all actions taking place for all extant and Extinct in the Wild species based on the IUCN classification scheme (Salafsky et al., 2008), including actions taking place to prepare for future reintroductions of Extinct in the Wild species. We used the 2019 version 3 IUCN Red List information in the plots for all species, including the two species that were Extinct in the Wild in 1993 (Przewalski's Horse *Equus ferus* and Guam Rail *Hypotaenidia owstoni*), and one species that was Extinct in the Wild in 2010 (Guam Rail *Hypotaenidia owstoni*), but which have been successfully reintroduced and are extant now (IUCN 2020).

We made these plots for those species with a median best probability > 50% that their extinction was prevented for the 1993 - 2020 time period (as shown in the Results), as well as for those species with a median best probability > 50% that their extinction was prevented for the 2010 - 2020 time period, and for all candidate species (Figures S4 - S10).

All code and data can be found at http://github.com/rbolam/Prevented_bird_and_mammal_extinctions.

### Extended results

#### Extinctions and Extinct in the Wild species

*Table S2. Bird and mammal species that have become extinct since 1993 (EX, Extinct), or are strongly suspected to have done so (CR(PE) – Critically Endangered (Possibly Extinct), i.e. species classified as Critically Endangered which are, on the balance of evidence, likely to be extinct, but for which there is a small chance that they may be extant).*

| **Species** | **2019 Red List category** | **Estimated date of extinction** |
|---|---|---|
| ***Birds*** | | |
| Maui Akepa<br>*Loxops ochraceus* | CR(PE) | 1994 |
| Least Vermilion Flycatcher<br>*Pyrocephalus dubius* | EX | 1994 |
| Imperial Woodpecker<br>*Campephilus imperialis* | CR(PE) | 1995 |
| Aguijan Reed-warbler<br>*Acrocephalus nijoi* | EX | 1996 |
| Glaucous Macaw<br>*Anodorhynchus glaucus* | CR(PE) | 2001 |
| Pernambuco Pygmy-owl<br>*Glaucidium mooreorum* | CR(PE) | 2001 |
| Poo-uli<br>*Melamprosops phaeosoma* | EX | 2004 |
| South Island Kokako<br>*Callaeas cinereus* | CR(PE) | 2007 |
| Cryptic Treehunter<br>*Cichlocolaptes mazarbarnetti* | EX | 2007 |
| Alagoas Foliage-gleaner<br>*Philydor novaesi* | EX | 2011 |
| ***Mammals*** | | |
| Telefomin Cuscus<br>*Phalanger matanim* | CR(PE) | 1998 |
| Yangtze River Dolphin<br>*Lipotes vexillifer* | CR(PE) | 2002 |
| Miss Waldron's Red Colobus<br>*Piliocolobus waldroni* | CR(PE) | 2008 |
| Christmas Island Pipistrelle<br>*Pipistrellus murrayi* | EX | 27 August 2009 |
| Bramble Cay Melomys<br>*Melomys rubicola* | EX | 2009 |

*Table S3. Number of species that went Extinct, for which extinction has been prevented, and the rate at which extinctions have been prevented.*

| | Birds | Mammals | Total |
|---|---|---|---|
| Extinctions between 1993-2020 | 10 | 5 | 15 |
| Extinctions between 2010-2020 | 1 | 0 | 1 |
| Species listed as Extinct in the Wild 1993-2020 | 6 | 3 | 8 |
| Species listed as Extinct in the Wild 2010-2020 | 6 | 2 | 7 |
| Species for which extinction is judged to have been likely in the absence of conservation during 1993-2020 (incl. Extinct in the Wild) | 21 – 32 | 7 – 16 | 28 – 48 |
| Species for which extinction is judged to have been likely in the absence of conservation during 2010-2020 (incl. Extinct in the Wild) | 9 – 18 | 2 – 7 | 11 – 25 |
| Ratio of prevented extinctions to extinctions 1993-2020 | 3.1 – 4.2 | 2.4 – 4.2 | 2.9 – 4.2 |
| Ratio of prevented extinctions to extinctions 2010-2020 | 10 – 19 | undefined | 12 – 26 |

*Table S4. Bird and mammal species that are currently Extinct in the Wild. All species are now held in ex situ collections only, with the exception of the reintroduced Przewalski's Horse and Guam Rail (see also comments in table).*

| **Species** | **Estimated date of extinction in the wild** |
|---|---|
| ***Birds*** | |
| Socorro Dove<br>*Zenaida graysoni* | 1972 |
| Guam Kingfisher<br>*Todiramphus cinnamominus* | 1986 |
| Guam Rail<br>*Hypotaenidia owstoni* | 1987 (this species was reintroduced from 2010, and was re-assessed as Critically Endangered in 2019). |
| Alagoas Curassow<br>*Mitu mitu* | Late 1980s |
| Spix's Macaw<br>*Cyanopsitta spixii* | 2000 |
| Hawaiian Crow<br>*Corvus hawaiiensis* | 2002 |
| ***Mammals*** | |
| Père David's Deer<br>*Elaphurus davidianus* | 1900 |
| Przewalski's Horse<br>*Equus ferus* | 1960s (this species was reintroduced from 1994, and was re-assessed as Critically Endangered in 2008. We considered it for time period 2010 - 2020) |
| Scimitar-horned Oryx<br>*Oryx dammah* | late 1980s-early 1990s |

*Table S5. Number of species by family, for all candidate species, species for which extinction is judged to have been likely to have occurred in the absence of conservation action during 1993-2020, and species for which extinction is judged to have been likely to have occurred in the absence of conservation action during 2010-2020. Ordered alphabetically. Number of candidate species: birds N = 45, mammals N = 24.*

| **Family** | **Candidate species** | **1993-2020 extinctions prevented** | **2010-2020 extinctions prevented** |
|---|---|---|---|
| *Birds* | | | |
| Acrocephalidae | 1 | 0 | 0 |
| Alcedinidae | 1 | 1 | 1 |
| Anatidae | 2 | 0 | 0 |
| Apterygidae | 1 | 0 | 0 |
| Callaeidae | 1 | 0 | 0 |
| Campephagidae | 1 | 1 | 1 |
| Cathartidae | 1 | 1 | 0 |
| Charadriidae | 1 | 1 | 1 |
| Columbidae | 2 | 2 | 1 |
| Corvidae | 2 | 2 | 2 |
| Cracidae | 2 | 2 | 1 |
| Laridae | 1 | 0 | 0 |
| Monarchidae | 3 | 3 | 2 |
| Muscicapidae | 1 | 1 | 0 |
| Otididae | 1 | 0 | 0 |
| Passerellidae | 2 | 2 | 0 |
| Procellariidae | 3 | 2 | 0 |
| Psittacidae | 9 | 7 | 4 |
| Rallidae | 1 | 1 | 1 |
| Recurvirostridae | 1 | 1 | 1 |
| Scolopacidae | 1 | 0 | 0 |
| Sturnidae | 1 | 1 | 1 |

| | | | |
|---|---|---|---|
| Thamnophilidae | 1 | 1 | 1 |
| Thraupidae | 1 | 1 | 1 |
| Threskiornithidae | 2 | 2 | 0 |
| Troglodytidae | 1 | 0 | 0 |
| Turdidae | 1 | 0 | 0 |
| *Mammals* | | | |
| Bovidae | 1 | 1 | 1 |
| Callitrichidae | 2 | 0 | 0 |
| Canidae | 1 | 1 | 1 |
| Cercopithecidae | 2 | 1 | 1 |
| Cervidae | 2 | 1 | 1 |
| Equidae | 1 | 1 | 0 |
| Felidae | 1 | 1 | 0 |
| Hylobatidae | 2 | 2 | 0 |
| Leporidae | 1 | 0 | 0 |
| Macropodidae | 2 | 0 | 0 |
| Mustelidae | 1 | 1 | 1 |
| Phocoenidae | 1 | 1 | 1 |
| Potoroidae | 1 | 1 | 0 |
| Rhinocerotidae | 2 | 1 | 1 |
| Sciuridae | 2 | 2 | 0 |
| Suidae | 1 | 1 | 0 |
| Vombatidae | 1 | 1 | 0 |

#### Further information for species for which extinction is judged to have been likely to have occurred in the absence of conservation action during 1993-2020

*Table S6. Species judged likely to have become Extinct in the Wild during 1993-2020 in the absence of conservation and for which captive populations exist.*

| **Species held in captivity** |
|---|
| *Birds* |
| Asian Crested Ibis *Nipponia nippon* |
| Bali Myna *Leucopsar rothschildi* |
| Black Stilt *Himantopus novaezelandiae* |
| California Condor *Gymnogyps californianus* |
| Echo Parakeet *Psittacula eques* |
| Lear's Macaw *Anodorhynchus leari* |
| Malherbe's Parakeet *Cyanoramphus malherbi* |
| Mariana Crow *Corvus kubaryi* |
| Northern Bald Ibis *Geronticus eremita* |
| Orange-bellied Parrot *Neophema chrysogaster* |
| Pink Pigeon *Nesoenas mayeri* |
| Puerto Rican Amazon *Amazona vittata* |
| Red-billed Curassow *Crax blumenbachii* |
| *Mammals* |
| Black-footed Ferret Mustela nigripes |
| Iberian Lynx *Lynx pardinus* |
| Przewalski's horse *Equus ferus* |
| Pygmy Hog *Porcula salvania* |
| Red Wolf *Canis rufus* |
| Vancouver Island Marmot *Marmota vancouverensis* |

*Table S7. Identified extant species and summary of key information. Birds: N = 26, mammals: N = 13.*

| Species | Medians for lowest, best and highest scores for 1993 - 2020 | Medians for lowest, best and highest scores for 2010 - 2020 | 1993 population estimate and trend | 2010 population estimate and trend | 2019 population estimate and trend | Threats | Conservation actions implemented | Why it is considered plausible that the species may have gone extinct |
|---|---|---|---|---|---|---|---|---|
| Alagoas Antwren *Myrmotherula snowi* | 80 - 91 - 99 | 70 - 85 - 95 | 16-160 mature individuals and a best estimate of 60, at four different sites, and declining. | 30 (range 10-100) mature individuals at one site remaining, and declining. | 12 individuals confirmed in 2019, and a range of 10-20 mature individuals, and declining. | Habitat loss through agriculture, housing, and logging | Protected Areas designated | Considering the small and declining population, if the Protected Areas had not protected some of the remnant vegetation, it is plausible that further habitat loss would have led to the extinction of the species. |
| Asian Crested Ibis *Nipponia nippon* | 80 - 90 - 98.5 | NA | 22 - 25 birds, and increasing. | 780 individuals, and c. 200 mature birds, and increasing. | At least c. 500 breeding pairs or 1000 mature birds in 2019, and increasing. | Habitat loss including wetlands and trees for nesting. Loss of food sources through agrochemical use, and conversion of rice paddies to wheat fields | Ban of agrochemical use, protection and guarding of nesting trees, maintenance of fields during winter for feeding, release of captive birds | The small number of individuals increased rapidly. The main threats were addressed directly, and plausibly led to the increase of the population. |

| | | | | | | | | |
|---|---|---|---|---|---|---|---|---|
| Bali Myna *Leucopsar rothschildi* | 85 - 95 - 100 | 80 - 90 - 98 | 42 - 48 individuals, and decreasing | 115 individuals but <50 mature individuals, and decreasing, as the population has been maintained only by release of captive birds. | 191 individuals in April 2019, with at least 100 birds released since October 2018, therefore no more than c.50 mature adults, and decreasing. | Illegal trapping for the cage-bird trade, and habitat loss | Release of captive individuals, Protected Area, legislation, sustainable livelihood projects | Considering the intensive conservation actions and the intensity of threat from illegal trapping, but the lack of increase of mature individuals in the population, it is plausible that the species would have gone extinct if not for the actions. |
| Black Stilt *Himantopus novaezelandiae* | 72.5 - 90 - 99 | 60 - 70 - 85 | 60 birds (estimate), and possibly slightly increasing. | 40 individuals, roughly equivalent to 27 mature individuals, and increasing | 106 adults in 2017, including released sub-adults and juveniles, therefore up to 49 mature individuals, and increaing. | Predation from both invasive and native species, and habitat loss through agriculture and hydroelectric developments, nest site destruction and disturbance, hybridisation with *Himantopus leucocephalus* | Captive rearing and release, with over 100 individuals released per year in recent years, predator exclusion fencing and trapping around nest sites, control of hybrids | Considering the intense threats to the species, the intensive management of the species by rearing chicks in captivity and then releasing them, and removal of predators around nests have plausibly prevented the extinction of the species. |
| California Condor *Gymnogyps californianus* | 92.5 - 98.5 - 100 | 32.5 - 50 - 70 | 4 individuals in the wild, | 104 adults in the wild, of which 44 have | 312 individuals in the wild in 3 | Lead poisoning, persecution, | Captive breeding and reintroduction | There were only four individuals in the wild in 1993, which were still |

| | | | | | | | | |
|---|---|---|---|---|---|---|---|---|
| | | | trend unknown. | produced viable offspring, and increasing. | meta-populations, and increasing | electrocution through powerlines, ingestion of plastics and other materials, thinning of eggshells due to DDT | programme, food provisioning, ban of lead ammunition and provision of lead-free ammunition, treatment for lead poisining in wild birds, protection of feeding habitat | threatened by lead poisoning, which is incremental. The release of further birds and intensive management both to reduce the use of lead ammunition, and to treat wild birds with lead poisoning, plausibly prevented the extinction of the species. |
| Chatham Island Taiko/Magenta Petrel *Pterodroma magentae* | 40 - 62.5 - 80 | 20 - 37.5 - 50 | A presumed total population of 45 - 70 or 100 - 150 individuals, trend unknown. | 18 known breeding pairs by 2012/2013 and 150-200 individuals estimated, trend stable or increasing (some non-genuine change) | In 2018/2019 114 adults were recorded in a season, resulting in a population estimate of 150-200 adults, and a genuine increase since 2014. | Predation of chicks and potentially adults by introduced species, habitat degradation through livestock, uneven sex ratios of adults | Control of invasives, protection of breeding areas, translocation | In 1993, there were only 4 known breeding pairs. While some of the increase in numbers is due to more burrows being found, there has also been a genuine increase since 2014. Considering the threat by invasives is managed intensively, it is plausible that extinction in this species has been prevented. |
| Echo Parakeet *Psittacula eques* | 82.5 - 94.5 - 99.5 | NA | 16 - 22 birds, including five pairs, and increasing. | 300-350 mature individuals estimated at the end of the 2011/2012 | No updated numbers. | Severe habitat loss (only 5% of native vegetation remained in 1995) leading | Captive breeding and release, intensive nest management including | The population increased from 16 - 22 birds in 1993 to 300 - 350 individuals at the end of the 2011/2012. Severe habitat loss and |

| | | | | | | | | |
|---|---|---|---|---|---|---|---|---|
| | | | | breeding season, and increasing rapidly. | | to loss of food sources and increased interspecific competition for nest sites, predation of chicks | provision of nest boxes and controlling nest predators, habitat management and restoration | predation are being addressed by habitat restoration, captive breeding and release, and nest site provision and protection, and it is plausible that these intensive actions have prevented extinction in this species. |
| Fatu Hiva Monarch *Pomarea whitneyi* | 75 - 90 - 98.5 | 75 - 90 - 96.5 | Numbers unknown, but persisting and common in 1990, and decreasing. | 64 individuals estimated in total population in 2009, and declining. | 31-36 individuals in 2019, and overall decreasing, but increasing since 2017. | Predation by invasives, some habitat loss | Intensive control of invasives | The rapid decline of the species was caused by invasive species, which are now being controlled and the species is increasing since 2017. It is plausible that the species would have gone extinct if not for the intensive predator control. |
| Guadalupe Junco *Junco insularis* | 75 - 90 - 96.5 | NA | 100 individuals (presumed), trend unknown. | Unknown, but thought to be less than 250 mature individuals, and thought to be increasing. | 10,900 - 39,800 individuals in 2018 and increasing | Decline in habitat through intensive grazing by goats, predation by invasives | Eradication of goats by 2007, control of cats | The species suffered from lack of habitat which was addressed by the eradication of goats, and its decline was exacerbated by predation, so it is plausible that the actions prevented extinction. |
| Lear's Macaw *Anodorhynchus leari* | 70 - 85 - 95 | NA | Around 60 - 100 individuals each in Raso | 1,123 birds in Raso da Catarina population | 1,700 birds in 2018 at Raso da Catarina, and | Illegal trapping and trade, deforestation, persecution for | Improved surveillance to stop illegal trapping and | It is plausible that the species would have been trapped to extinction given one of the |

| | | | | | | | | |
|---|---|---|---|---|---|---|---|---|
| | | | da Catarina and Boqueirão da Onça populations, one of which was increasing, the other decreasing. | which is increasing, 2 in the Boqueirão da Onça population | increasing. 2 in the Boqueirão da Onça population | foraging on maize crops | trade, maize replacement scheme for farmers | populations declined from 60 - 100 individuals to 2, if not for the actions to stop poaching and smuggling |
| Mangrove Finch *Geospiza heliobates* | 60 - 70 - 90 | 40 - 60 - 70 | 100 - 200 birds, presumed stable | 80-120 mature individuals, and decreasing | 41 birds were observed, and the population estimate is 80 – 100, and is decreasing | Disease, climate change, nest predation | Protected Area, control of invasives, captive rearing of young, treatment of nests to reduce number of parasites | The population has declined over the time period, and the species is intensively managed to reduce predation and disease caused by nest parasites. This plausibly prevented extinction in this species. |
| Mariana Crow *Corvus kubaryi* | 60 - 80 - 95 | 50 - 70 - 85 | 891 individuals on Rota and less than 50 individuals on Guam, and declining rapidly | 60 confirmed pairs on Rota in 2008, two males on Guam, and decreasing rapidly | 50 breeding pairs in the 2015-2016 breeding season, and decreasing | Predation by invasives, habitat loss, direct persecution | Control of invasives, screening to prevent invasives becoming established | The species went Extinct on Guam due to invasive Brown Tree Snakes, and prevention of these snakes to become established on Rota plausibly prevented extinction in this species. |
| Northern Bald Ibis *Geronticus eremita* | 40 - 65.5 - 80 | 17.5 - 30 - 50 | 59 pairs in 1997 following the death of 40 birds in 1996 | 105 pairs, but only four mature birds in Syria in 2009, overall stable | 708 individuals as of 2018, and increasing | Disturbance, agricultural intensification, hunting, poisoning | Protected Areas, community involvement to prevent | Considering the small population at the beginning of the period, it is plausible that the Protected Area and |

| | | | | | | | | |
|---|---|---|---|---|---|---|---|---|
| | | | | | | | disturbance, water provisioning, reintroduction | community involvement have mitigated threats such as habitat loss and disturbance, and prevented extinction. |
| Orange-bellied Parrot *Neophema chrysogaster* | 85 - 95 - 99 | 80 - 90 - 98.5 | 150 estimated, possibly stable | <50 individuals, and decreasing | Total population was 14 in 2017, and decreasing | Habitat loss, disease, competition and predation | Protection and management of feeding habitat, release of large numbers of captive individuals (partly successful) | Only 14 individuals were remaining as of 2017, after a steady decline. It is plausible that the actions have slowed the decline enough to prevent extinction by 2020. |
| Orange-fronted Parakeet; Kakariki Karaka; Malherbe's Parakeet *Cyanoramphus malherbi* | 50 - 80 - 90 | 40 - 65 - 80 | Estimate of 150 – 200 individuals in 1999, and likely to have been declining | 450 individuals but uncertainty around the number of mature individuals. Increasing | 380 individuals, and increasing | Invasive species, habitat alteration | Intensive control of invasives, nest site protection, translocation | The species was declining in the early 2000s mainly through predation by invasives. Successful control of these, as well as translocations (one of which was sucessful) mean it is plausible that extinction was prevented. |
| Pale-headed Brush-finch *Atlapetes pallidiceps* | 75 - 85 - 95 | NA | 12 - 22 occupied territories in 1999 estimate, and decreasing | 226 mature individuals, and increasing | Estimate of 90-104 territories, and stable | Brood parasitism, lack of suitbale habitat | Habitat protection, removal of brood parasites, habitat management | The small population increased rapidly once brood parasites were being removed, which plausibly prevented extinction in this species. |

| | | | | | | | | |
|---|---|---|---|---|---|---|---|---|
| Pink Pigeon *Nesoenas mayeri* | 90 - 95 - 100 | NA | Wild population of 20, and introduced population of 28, and increasing | 360-395 individuals in 2005, and roughly stable (fluctuating) | Known population of c.325 to c.410 individuals, and stable | Habitat loss, predation by invasives | Protected Area, habitat restoration, nest protection and control of invasive predators, supplementary feeding, captive breeding and release | The small population increased rapidly due to intensive conservation effforts which plausibly prevented extinction. |
| Puerto Rican Amazon *Amazona vittata* | 90 - 97 - 100 | 50 - 70 - 85 | 41 birds, and ncreasing | 50-70 individuals in 2 populations, roughly equivalent to 33-47 mature individuals, and increasing | 70-80 wild parrots in the reintroduced population, the original population disappeared following a hurricane. Increasing | Habitat destruction by hurricanes, competition, predation (by native and invasive species), parasitism of chicks | Nest site and food provision, control of nest predators and competitors, captive breeding and release, habitat protection | The original population went extinct following a hurricane in 2017, leaving only the reintroduced population, and many other conservation actions are addressing threats directly. It is therefore plausible that actions have prevented its extinction. |
| Rarotonga Monarch *Pomarea dimidiata* | 75 - 90 - 95 | NA | 60 individuals, and increasing | 380 birds estimated in 2011, and increasing | 500 mature individuals estimated, and increasing | Invasive species | Intensive control of invasives, translocation | The population has been increasing. The key threat of invasives is being addressed by intensive control during the breeding season, and it is plausible this prevented extinction in this species. |

| | | | | | | | | |
|---|---|---|---|---|---|---|---|---|
| Red-billed Curassow *Crax blumenbachii* | 35 - 60 - 80 | NA | No population estimates, but considered very small and possibly decreasing | Around 500 native individuals, and possibly decreasing | Around 500 individuals, and different estimates for trends | Habitat loss, hunting | Protected Areas, reintroductions | Habitat loss is a key threat to this species, exacerbated by hunting, and the species is now laregly restricted to actively protected reserves. It is therefore plausible that extinction has been prevented. |
| Reunion Cuckooshrike *Lalage newtoni* | 60 - 76.5 - 90 | 30 - 57.5 - 75 | 120 pairs, and stable | 30 pairs and decreasing | 33 pairs in 2013, and decreasing | Predation and competition with invasives, poaching, disease, habitat loss | Habitat protection and management, control of invasives, control of hunting, curbing of tourism | The species has been declining, and suffers from a wide range of threats. Intensive conservation actions have plausibly prevented its extinction. |
| Seychelles Magpie-robin *Copsychus sechellarum* | 77.5 - 90 - 99 | 15 - 39 - 53 | 46 birds on one island in 1994, and increasing | 207 individuals on five islands, and increasing | 283 birds on 5 islands in 2015, and increasing | Invasives, predation, pesticide use | Translocations to predator-free islands, control of invasives, ban of pesticide use | The population increased and due to translocation the species now exists on five rather than one island. In addirion, invasives were controlled and pesticides banned, plausibly preventing extinction in this species. |
| Southern Red-breasted Plover *Charadrius obscurus* | 75 - 90 - 96.5 | 50 - 65 - 80 | 60-65 individuals, and declining | Estimated at 288 individuals, and fluctuating | Estimated at 170 individuals. Decreasing rapidly between | Invasive species | Control of invasives, which intensified after species decreased | The species went extinct on one island due to invasive species, which are a threat elsewhere too. Intensive control of invasives make it plausibe |

| | | | | | | | | |
|---|---|---|---|---|---|---|---|---|
| | | | | | 2012 - 2016, but now increasing. | | rapidly since 2012 | that extinction has been prevented. |
| Tahiti Monarch *Pomarea nigra* | 77.5 - 90 - 97.5 | 60 - 80 - 90 | Several pairs in 4 different valleys, trend unknown | 35 individuals, and increasing | 79 mature individuals, and increasing | Habitat loss and invasive species, which are causing habitat changes and predation | Control of invasive plants and animals, and planting of food plants | The species had a small population that was facing many threats, including habitat loss and invasive species. Actions have addressed these threats, particularly the invasives, and the species is now increasing. Therefore extinction was plausibly prevented. |
| Yellow-eared Parrot *Ognorhynchus icterotis* | 62.5 - 80 - 90 | NA | Few records of this species, and declining | In Colombia, 1,103 individuals including 106 adult pairs and increasing. In Ecuador, the last individuals were reported in 1998. | 2,601 individuals in 2019, and increasing | Habitat loss, especially loss of wax palms as the main habitat for this species, hunting | Habitat protection and restoration, awareness campaign to stop the use of wax palm, provision of nest boxes | The species recovered from just a few individuals to over 2,000 by 2019. There was little habitat remaining, and the species was hunted, but habitat protection and restoration have been very successful alongside a public awareness campaign and ban of the use of wax palms. The actions plausibly prevented extinction in this species. |
| Zino's Petrel *Pterodroma madeira* | 25 - 57.5 - 75 | 12.5 - 30 - 50 | 20 - 30 known pairs, and stable | 130-160 individuals estimated, and stable | 200 individuals, and stable | Predation by invasives (on one occastion | Control of invsive species | The population increase is partly due to increased search effort, but some genuine change is also |

| | | | | | | | | |
|---|---|---|---|---|---|---|---|---|
| | | | | | | a cat killed 10 birds), fire | | recorded. Invasive species are the main threat and have been controlled, plausibly preventing extinction. |
| ***Mammals*** | | | | | | | | |
| Black-footed Ferret *Mustela nigripes* | 90 - 95 - 100 | 40 - 60 - 72.5 | The species was Extinct in the Wild, and by 1993 there were approximatel y 10-20 reintroduced ferrets. Trend unknown as species was only reintroduced 2 years previously | 448 breeding adults in 2009 which declined to 274 in 2012 | 112 breeding adults recorded in 2019, but due to incomplete survey efforts actual number is probably closer to 240, and decreasing | Disease which affected both ferrets as well as their main prey base, risk of inbreeding, lack of suitable habitat | Ongoing release of captive-bred individuals | The species was reintroduced just prior to 1993, and through ongoing releases first increased, but has been decreasing again since 2009. There are substantial reintroduction efforts ongoing, with 148 kits released in 2019 for example, plausibly preveting extinction in this species. |
| Cao-vit Gibbon *Nomascus nasutus* | 40 - 60 - 80 | 20 - 37.5 - 60 | This species was thought to be possibly extinct, but a surviving population was found in 2002 | 2005 population estimate of 35-37 individuals in Vietnam, and at least 10 individuals found in China in 2006, and decreasing | Overall population of 129, trend unknown | Habitat loss for charcoal, grazing, and cultivation; hunting | Habitat conservation and reduction of charcoal use, patrols to stop hunting | The species persisted until 2002, when it was rediscovered. Actions have addressed the main threats by protecting habitat and patrolling to stop hunting, hence extinction has plausibly been prevented. |
| Cat Ba Langur | 60 - 75 - 90 | 40 - 55 - 75 | 104 to 135 individuals in | Estimate of 50, trend unknown | Total population of | Poaching led to severe declines | Controls to stop poachers, | The species declined rapidly due to poaching, |

| | | | | | | | | |
|---|---|---|---|---|---|---|---|---|
| *Trachypithecus poliocephalus* | | | 1999/2000, and declining | | 67 individuals, and approximately 35 mature individuals, trend unknown | and small population is at risk of inbreeding effects; habitat destruction | protected areas | which is now being controlled. It occurs in Protected Areas. Therefore extinction has plausibly been prevented. |
| Gilbert's Potoroo *Potorous gilbertii* | 95 - 100 - 100 | 25 - 40 - 65 | 30 or less in 1999 in one population, trend unknown | 60 - 100 mature individuals in two subpopulations, and stable | 45 individuals in one population. Stable as of 2019, but delining previously | Predation by introduced species, risk of fire which led to extinction of original population in 2015 | Predator control, translocation onto predator-free island | This species has had low numbers, and the threat of fire is so severe that the original population went extinct. A translocated population was established prior to 2010 and remains, plausibly preventing extinction in this species. |
| Hainan Gibbon *Nomascus hainanus* | 50 - 80 - 95 | 30 - 50 - 70 | Three groups with less than 20 individuals, and decreasing | 20 individuals in two groups, with some solitary individuals, and stable | More than 25 individuals, and stable | Hunting, lack of suitable habitat, small population size | Entire range within a Protected Area | This species has a small population size, which seems to be stable. It is threatened by poaching and lack of habitat. As its entire range is within a protected area, it is plausible that this prevented its extinction. |
| Iberian Lynx *Lynx pardinus* | 50 - 80 - 90 | 20 - 32.5 - 50 | 725 mature individuals in 1985, and 65 mature individuals in | Estimate of 130, and increasing | Estimate of 320 for 2018, and increasing | Strong dependence on rabbit as a prey base which had declined due | Actions to boost rabbit numbers, reduce road casualties, education to | This species declined rapidly and is facing many threats, which are being tackled comprehensively. It is now increasing, and it is plausible that |

| | | | | | | | | |
|---|---|---|---|---|---|---|---|---|
| | | | 2001, hence decreasing | | | to disease, shooting and trapping, road casualties, lack of habitat, habitat loss, small populations showing poor reproduction and genetic performance | stop trapping, translocations to stop inbreeding and many reintroduction projects, protected areas | extinction has been prevented. |
| Javan Rhinoceros *Rhinoceros sondaicus* | 80 - 90 - 100 | 50 - 75 - 87.5 | 35 - 50, and possibly stable | 40 - 60, and possibly stable | A minimum of 67, and stable | Poaching | Listed on CITES, protected areas, rhino protection units to stop poachers | The severe threat from poaching means that it is plausible the small population could have been hunted to extinction, if not for efforts to stop poachers. |
| Northern Hairy-nosed Wombat *Lasiorhinus krefftii* | 50 - 80 - 95 | 20 - 32.5 - 55 | Estimate of 65, and increasing | Estimate of 162, and increasing | 2016 estimate of 245, and probably increasing | Competition with grazing animals, predation, inbreeding | Fence to exclude predators, translocation to establish a second population, cutting of introduced flora to promote growth of native flora, | The species faces different threats which are being addressed through intensive management such as fences to exclude predators, a successfu translocation to establish a 2nd population, and removal of invasive plants to increase native vegetation, hence it is plausible that extinction has been prevented. |

| | | | | | | | | |
|---|---|---|---|---|---|---|---|---|
| | | | | | | | water provisioning | |
| Northern Idaho Ground Squirrel *Urocitellus brunneus* | 30 - 55 - 70 | NA | 5,000 individuals in 1985 which decreased to 450 to 500 individuals at 22 sites in 2002 | 1,560 across 56 sites, and increasing | 2,659 individuals, and less than 1,000 breeding adults, in 2016, and increasing | Habitat loss and fragmentation, competition, shooting and poisoning | Habitat management, regulatory changes that may have reduced the threat of shooting and poisoning. | The species was declining rapidly, and the threat of habitat loss has been managed since, plausibly preventing extinction in this species. |
| Pygmy Hog *Porcula salvania* | 50 - 70 - 80 | 20 - 40 - 60 | In the mid-1990s the population was between 400 and 500 individuals, and declining | The total population may have been ca. 300 in the wild, and stable | 100 hogs at release site, and stable (presumably original population is persisting) | Habitat loss through agriculture, forestry, human settlements, and flood control | Habitat protection, translocation | The severe pressure on habitats led to a loss of some populations prior to 1993, and could plausibly have driven the species to extinction if not for habitat protection efforts. |
| Red Wolf *Canis rufus* | 65 - 85 - 96.5 | 40 - 60 - 80 | 50, and increasing | more than 150 animals in 2005, trend in 2010 decreasing | Now restricted to federal lands, with 20 - 30 individuals, and declining | Hybridisation with coyotes, illegal killing | Reintroduction, Protected Areas | The species was reintroduced prior to 1993, and it increased in number until 2005. Threats are hybridisation with coyotes, and illegal killing of wolves which has increased as a result of conflicts with landowners. The species is now only protected in three wildlife refuges. Extinction was plausibly prevented. |

| | | | | | | | | |
|---|---|---|---|---|---|---|---|---|
| Vancouver Island Marmot *Marmota vancouverensis* | 50 - 72.5 - 90 | 20 - 45 - 60 | Estimated 300-350 individuals in 1986, followed by precipitous decline and near-extinction in the wild by 2000 | In 2007 it was estimated that there were about 85 individuals remaining in the wild, and decreasing | 140-190 in 2017, based on field counts, and decreasing | Ecosystem modification, native predators and invasives | Captive breeding programme and releases | This species appears to fluctuate dramatically. The population has been reinforced by releases of captive bred marmots, resulting in now two meta-populations, which plausibly prevented extinction. |
| Vaquita *Phocoena sinus* | 65 - 90 - 100 | 50 - 77.5 - 100 | In 1997 abundance was estimated to be 567, and declining | 245, and declining | 6 individuals recorded in summer 2018, and population estimate of 10 - 22, and declining | Illegal fishing for Totoaba causes the species to get tangled in nets and die | Ban on fishing totoaba, removal of illegal fishing gear, provision of alternative livelihoods | The species rapidly declined due to intensive fishing pressure in which it dies as bycatch. Despite bans and removal of fishing gear, fishing is ongoing. It is plausible that the actions have slowed the decline somewhat, and therefore prevented extinction by 2020. |

#### Plots for species for which extinction is judged to have been likely to have occurred in the absence of conservation action during 2010-2020

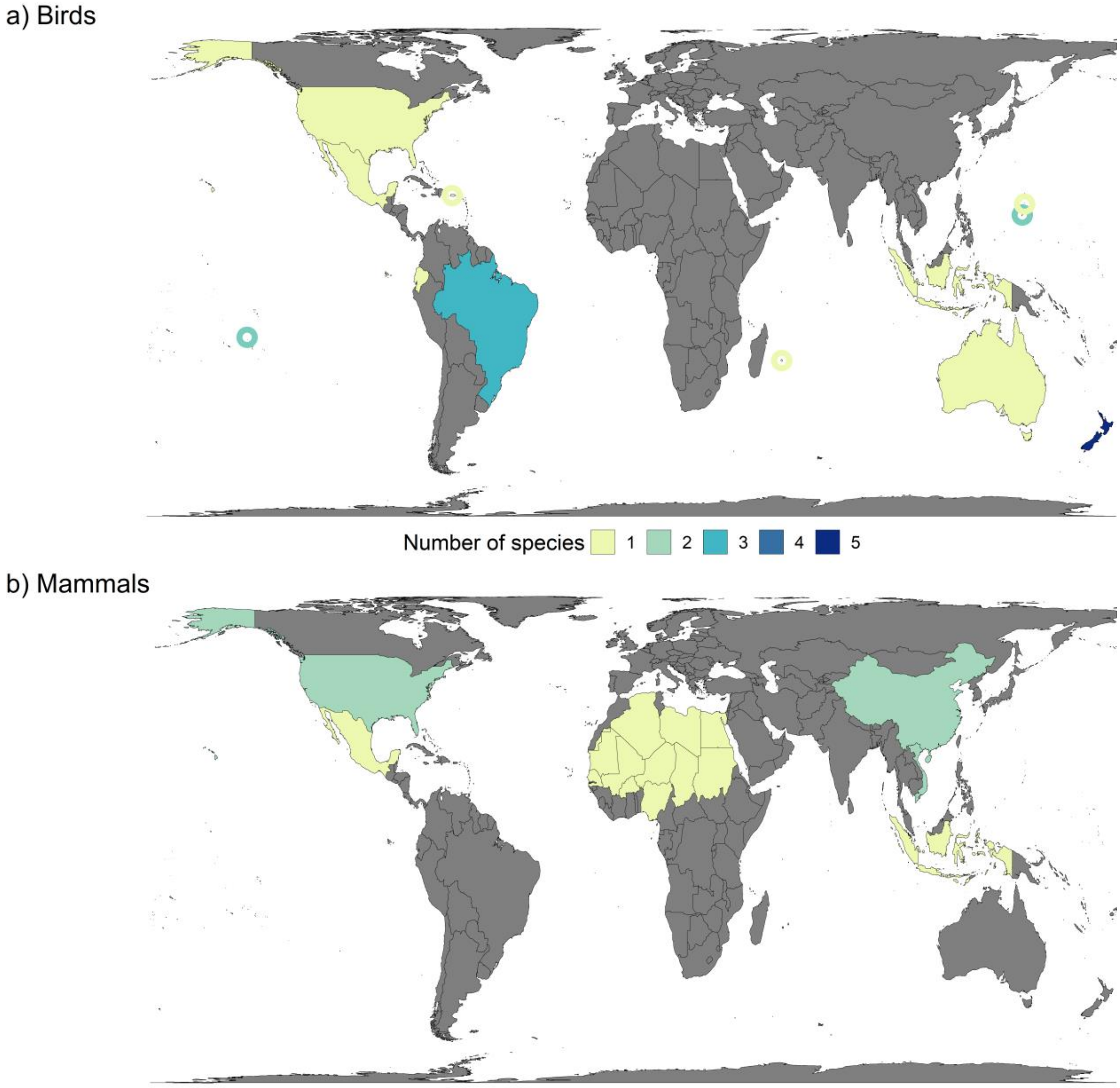


*Figure S1. Number of (a) bird (N = 18) and (b) mammal (N = 7) species for which extinction is likely to have occurred (i.e. median probability >50%) in the absence of conservation action during 2010-2020, per country. Circles show small island nations and overseas territories, and are coloured according to the key. Species listed as Extinct in the Wild (IUCN, 2020) were mapped in the last countries where they occurred, or are presumed to have occurred.*

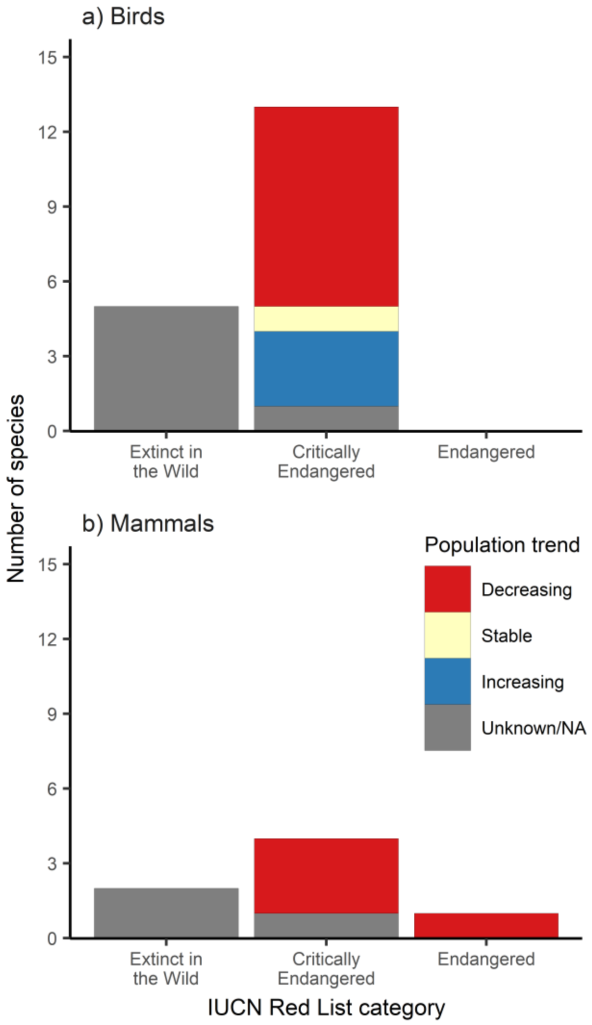


*Figure S2. 2019 IUCN Red List categories and population trends of (a) bird (N = 18) and (b) mammal (N = 7) species for which extinction is judged to have been likely (i.e. median probability >50%) to have occurred in the absence of conservation action, during 2010-2020.*

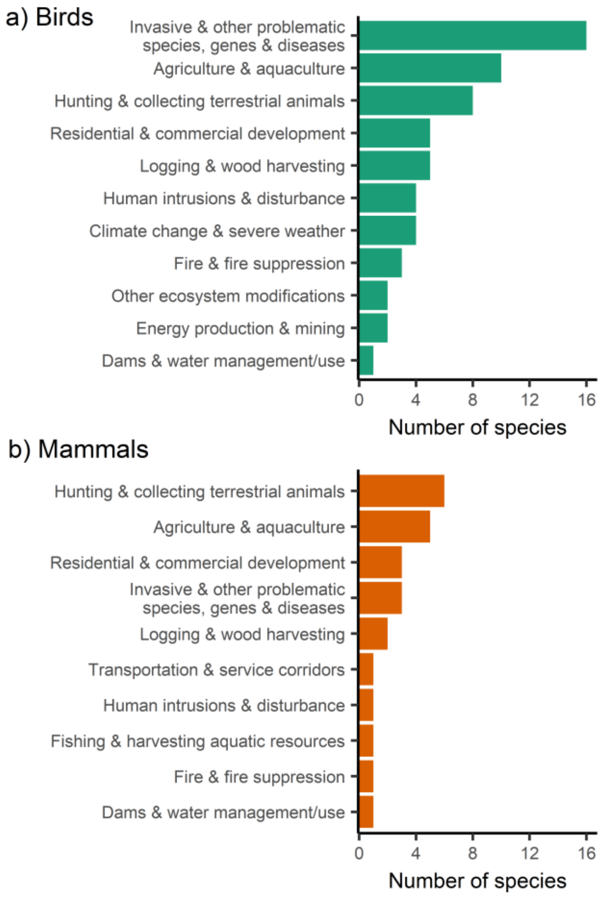


*Figure S3. Current and past threats to (a) bird (N = 18) and (b) mammal (N = 7) species for which extinction is judged to have been likely (i.e. median probability >50%) to have occurred in the absence of conservation action during 2010-2020. Threats are taken from the IUCN threat classification scheme level 1 (Salafsky et al., 2008).*

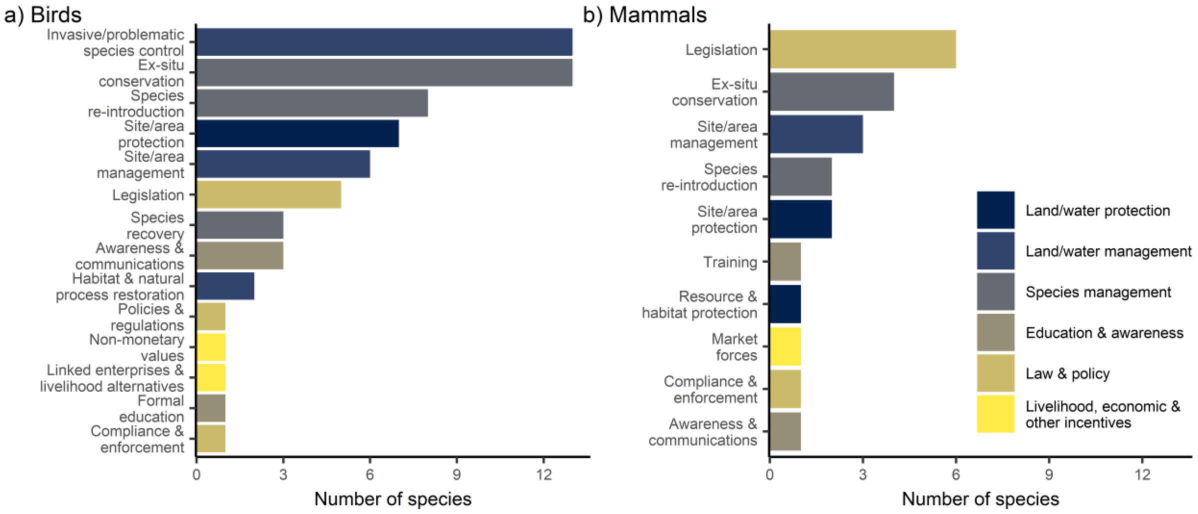


*Figure S4. Conservation actions for (a) bird (N = 18 and (b) mammal (N = 7) species for which extinction is judged to have been likely (i.e. median probability >50%) to have occurred in the absence of conservation action during 2010-2020. Actions are taken from the IUCN action classification scheme level 2, while colours denote level 1 (Salafsky et al., 2008). Both in-situ and ex-situ actions are included for species that are Extinct in the Wild.*

#### Plots for candidate species

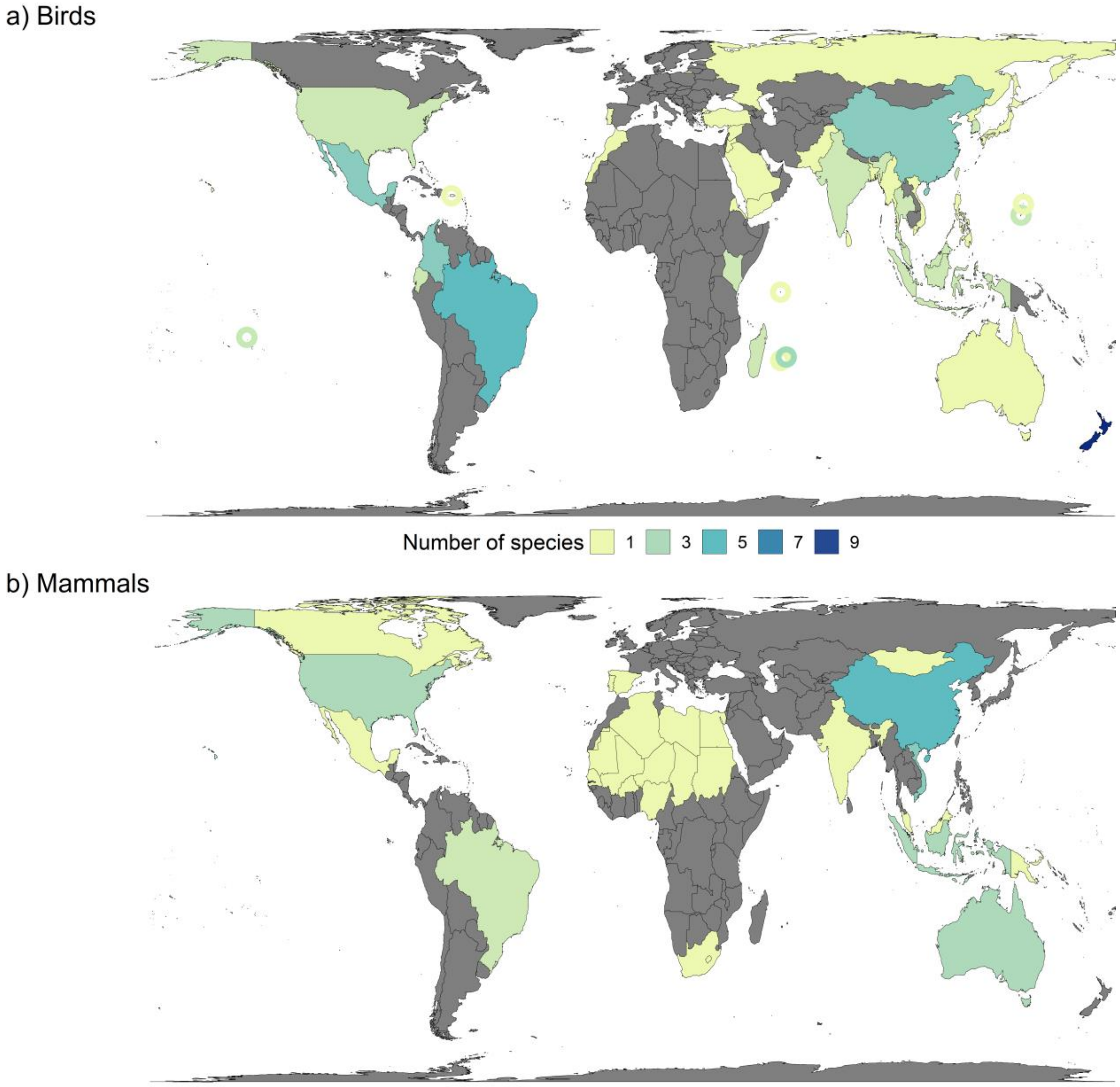


*Figure S5. Number of (a) bird (N = 45) and (b) mammal (N = 24) candidate species per country. Circles show small island nations and overseas territories, and are coloured according to the key. Species listed as Extinct in the Wild (IUCN, 2020) were mapped in the last countries where they occurred, or are presumed to have occurred.*

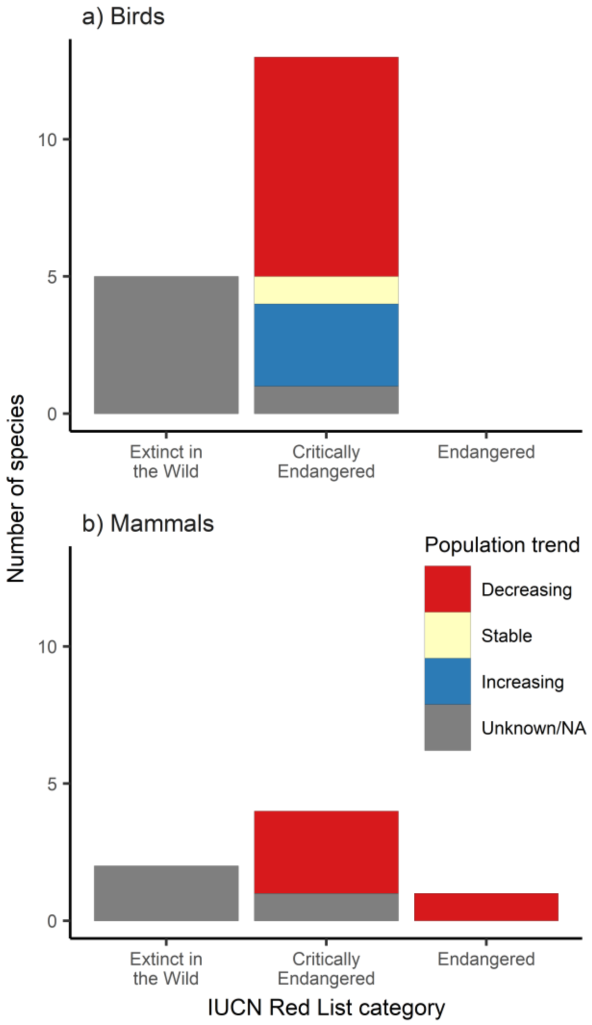


*Figure S6. 2019 IUCN Red List categories and population trends of (a) bird (N = 45) and (b) mammal (N = 24) candidate species.*

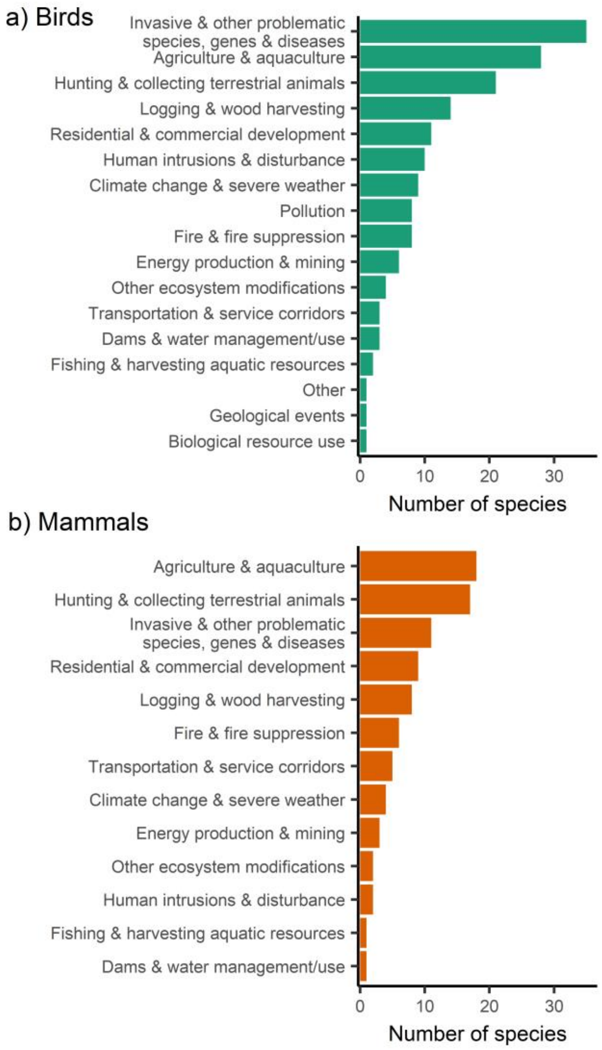


*Figure S7. Current and past threats to (a) bird (N = 45) and (b) mammal (N = 24) candidate species. Threats are taken from the IUCN threat classification scheme level 1 (Salafsky et al., 2008).*

#### Comparison between calls

We compared the scores between the two conference calls for each species and each time period. Species of note are those where the overall median best estimate is different to the median best estimate from the second call, because new information gained during the first call was available to evaluators on the second call, but not vice versa.

Of the 39 birds for 1993 - 2020, there were significant differences in the best estimates for Alagoas Antwren, Chatham Parakeet, Taita Thrush, and Yellow-eared Parrot. This difference would have resulted in exclusion if only the second call had been considered for only one of those species: **Yellow-eared Parrot**, with an overall median of 80%, a median of 90% for the first call, and of 25% for the second call. An additional species with no significant difference, but for which inclusion changed, was **Zino's Petrel**, with an overall median of 57.5%, a median of 65% for the first call, and of 45% for the second call. Of the 17 birds for 2010 - 2020, there were significant differences only for Townsend's Shearwater, but no impacts in which species scored >50%.

Of the 22 mammals for 1993 - 2020, there were significant differences for Cat Ba Langur, Riverine Rabbit, and Vancouver Island Marmot. A species with no significant difference, but for which inclusion changed, was **Delacour's Langur**, with an overall median of 50%, a median of 40% for the first call, and of 60% for the second call. Of the 17 mammals for 2010 - 2020, there were significant differences for Cat Ba Langur, Northern Hairy-nosed Wombat, Pygmy Hog, Tenkile, and Vancouver Island Marmot. The inclusion only changed for **Pygmy Hog** for 2010 - 2020, with an overall median of 40%, a median of 20% for the first call, and of 52.5% for the second call. An additional species with no significant difference, but for which inclusion changed, was **Red Wolf**, with an overall median of 60%, a median of 75% for the first call, and of 50% for the second call.

#### Information gained during or after the second calls

##### Yellow-eared Parrot

The Yellow-Eared Parrot population at Roncesvalles in the Western Andes has received conservation action, and was the only known population up until 2009. During the second conference call, a further population of Yellow-Eared Parrot in the Eastern Andes was mentioned (Murcia-Nova et al., 2009), and the possibility that the species may have persisted elsewhere in its range in the Western Andes (Renjifo et al., 2014), which led evaluators to give lower scores compared with the first call. There were no direct management actions taking place for these populations at the time of their discovery, but an awareness campaign taking place nationally may have also benefitted those populations by reducing habitat loss and hunting. By 2009, the Roncesvalles population had increased in size and small flocks would start to recolonise other areas. It is therefore possible that the populations in the Western Andes originate from the population at Roncesvalles, though there is no direct evidence for this. There are different opinions as to the origin of the population in the Eastern Andes. In the original description of when the population was found, it is mentioned that local people have known about this parrot population since the 1960s (Murcia-Nova et al., 2009), and another source details that this population also differs in the type of palms used by the species (Renjifo et al., 2014). Another source states that it is possible for this population to originate from the Roncesvalles population, which is 170km apart (Salaman et al., 2019).

##### Zino's Petrel

One evaluator argued that it is possible this species would have persisted without action. There was some non-genuine change recorded as the searching efforts for nest sites were increased. Some nests may have been successful without conservation as invasive species may not have been able to get to all of the petrel nests due to the inaccessible nature of its preferred nest sites.

##### Pygmy Hog

It was mentioned during the second call that the habitat of this species is under intense pressure. While conservation actions in the 1990s were ineffective, action was ramped up in the early 2000s, without which it is possible that all remaining habitat would have been lost.

##### Red Wolf

On the first mammal call, the current population of Red Wolf was understood to include only those individuals reintroduced in North Carolina. However, during the second call, it was mentioned that a second population of Red Wolf survived. One of the evaluators conveyed that Red Wolf DNA had been found in a pack of wolves on Galveston Island, an unprotected island off the coast of Texas. Although this population is clearly admixed (coyote and Red Wolf), genetic results suggest the animals are more close to captive Red Wolves than south-eastern coyotes (Heppenheimer et al., 2018 ). This introduced some uncertainty as to whether or not Red Wolf would have gone extinct in the absence of conservation, with this uncertainty increased due to the taxonomic ambivalent status of this population. It was noted that Red Wolves themselves have an ambivalent taxonomic status, but recent evidence has concluded that the Red Wolf should be considered a distinct species from the Grey Wolf and Coyote with likely historical admixture (National Academies of Sciences, Engineering, and Medicine 2019).

#### Comparison of results

Some of our results differ from those of Hoffmann et al. (2015), who estimated change in extinction risk for the world's ungulates in a scenario where all conservation ended between 1996 and 2008. Both studies consider that *Rhinoceros sondaicus* and *Equus ferus* would have become extinct (Extinct or Extinct in the Wild). However, whereas Hoffmann et al. assigned a category of Extinct as the best estimate for *Axis kuhlii* and *Rhinoceros unicornis*, the current study assigns them a lower probability of extinction due to new information that has become available since 2015. Finally, Arabian Oryx *Oryx leucoryx* was considered to have gone extinct by Hoffmann et al., (2015), but was excluded in our exercise during the process of identifying the candidate list of mammals. The 2003 Red List account of this species details that there were 886 individuals in 5 populations, which had increased to 1,220 individuals as of 2017 (850 mature individuals), and that these populations were stable or decreasing (IUCN SSC Antelope Specialist Group, 2017), which is why it was excluded. However, considering the findings from Hoffmann et al. (2015) we recognise that this species should have been included in our expert elicitation exercise.

Some of our results also differ from Young et al. (2014) who considered the counterfactual Red List assessment if actions had ceased for the time period 1988 - 2012. Both their and our results indicate that extinction or extinction in the wild has been prevented for Pink Pigeon *Nesoenas mayeri* and Echo Parakeet *Psittacula eques*. However, they also considered the extinction of Rodrigues warbler *Acrocephalus rodericanus* to have been avoided, whereas this species was given a median probability <50% in our Delphi exercise. We did not include Mauritius Kestrel *Falco punctatus* in our analysis, because most actions ceased in 1994 when the population was at 222-286 individuals (BirdLife International 2016). We did not consider Rodrigues Fody *Foudia flavicans* as it was at worst Vulnerable for the time period under consideration. Its population was increasing in 1993, and presumably between the 1983 estimate of 110 birds and 1999 estimate of 334 - 500 pairs (911 - 1,200 individuals).

### Additional references


BirdLife International (2016). Falco punctatus. The IUCN Red List of Threatened Species 2016: e.T22696373A93557909. https://dx.doi.org/10.2305/IUCN.UK.2016-3.RLTS.T22696373A93557909.en Downloaded on 05 February 2020.

Butchart, S. H. M., Akçakaya, H. R., Chanson, J., Baillie, J. E. M., Collen, B., Quader, S., ... Mace, G. M. (2007) Improvements to the Red List Index. PLoS ONE 2: e140. doi: 10.1371/journal.pone.0000140

Diamond, I.R., Grant, R.C., Feldman, B.M., Pencharz,P.B., Ling, S.C., Moore, A.M. & Wales, P.W. (2014). Defining consensus: a systematic review recommends methodologic criteria for reporting of Delphi studies. Journal of Clinical Epidemiology, 67, 401–409. doi: 10.1016/j.jclinepi.2013.12.002

Heppenheimer, E., Brzeski, K.E., Wooten, R., Waddell, W., Rutledge, L.Y., Chamberlain, M.J., Stahler, D.R., Hinton, J.W. and VonHoldt, B.M. (2018). Rediscovery of red wolf ghost alleles in a canid population along the American Gulf Coast. *Genes*, *9*(12), p.618.

Hoffmann, M., Belant, J.L., Chanson, J.S., Cox, N.A., Lamoreux, J., Rodrigues, A.S., … Stuart, S.N. (2011). The changing fates of the world's mammals. *Philosophical Transactions of the Royal Society B: Biological Sciences, 366*(1578), 2598-2610. *doi:* 10.1098/rstb.2011.0116

IUCN SSC Antelope Specialist Group (2017). Oryx leucoryx. The IUCN Red List of Threatened Species 2017: e.T15569A50191626. https://dx.doi.org/10.2305/IUCN.UK.2017-2.RLTS.T15569A50191626.en. Downloaded on 10 February 2020

Knol, A.B., Slottje, P., van der Sluijs, J.P. and Lebret, E. (2010). The use of expert elicitation in environmental health impact assessment: a seven step procedure. Environmental Health, 9(1), 19. doi: 10.1186/1476-069X-9-19

Murcia-Nova, M.A., Beltrán-Alvarado, D. and Carvajal–Rojas, L. (2009). Un nuevo registro del loro orejiamarillo (Ognorhynchus icterotis: Psittacidae) en la cordillera Oriental colombiana. *Ornitología Colombiana*, *8*, 94-99.

National Academies of Sciences, Engineering, and Medicine (2019). Evaluating the Taxonomic Status of the Mexican Gray Wolf and the Red Wolf. Washington, DC: The National Academies Press. doi: https://doi.org/10.17226/25351

Renjifo, L. M., Gómez, M. F., Velásquez-Tibatá, J., Amaya-Villarreal, A. M., Kattan, G. H., Amaya-Espinel, J. D., y Burbano- Girón, J. (2014). Libro rojo de aves de Colombia, Volumen I: bosques húmedos de los Andes y la costa Pacífica. Editorial Pontificia Universidad Javeriana e Instituto Alexander von Humboldt. Bogotá D.C., Colombia.

Salaman, P., Cortes, A., Waught, D. (2019) Back from the brink of extinction: how the recovery of the Yellow-Eared Parrot united a nation. Conservacion colombiana, 26. Available at https://www.proaves.org/wp-content/uploads/2020/01/Conservacion_Colombiana_26_21-35-Loros-Salaman-et-al.pdf (last accessed 05/02/2020)

### Information circulated to evaluators

**Project aim**

This project aims to quantify the number of mammal and bird extinctions that have been prevented by conservation action since 1993 (the inception of the Convention on Biological Diversity) and since 2010 (the adoption of Aichi Target 12, which aimed to prevent "the extinction of known threatened species"). The work is of great interest to the CBD Secretariat and Parties who are meeting in 2020 to launch the Global Biodiversity Outlook 5 (reviewing achievement of the Aichi Targets) and to adopt a new post-2020 biodiversity framework and targets.

**What will this exercise entail?**

If you agree to participate, you will be asked to review information we have compiled in a standard format for 23 mammal species that are candidates for having their extinction prevented by conservation action during one or both time periods. You will be asked to estimate confidentially the probability that the species would have gone extinct without action for one or both of the time periods (**Round 1**). We estimate this will take 4 – 6 hours, though it will vary from person to person and also depend on how much time you spend seeking additional information. You will then be asked to join a conference call to briefly discuss each species (considering the median and range of scores across the group) and then revise your scores confidentially (**Round 2**). For the conference call, you will be asked beforehand to familiarise yourself with the information for a particular small selection of the species (to be provided beforehand) to contribute especially to the discussion during the conference call. All scores will be anonymous.

Except for the conference call, all other tasks can be completed in your own time within the designated timeframes. You will be asked not to discuss the scores or other details with anyone else who is also involved in this elicitation except during the conference call.

An overview of the timeline for this exercise can be found in Table 1.

*Table 1. Key dates for expert elicitation exercise and time requirement.*

| Dates | Task | Time required |
|---|---|---|
| Before 20 December | **Round 1** - Score probabilities in spreadsheet that will be provided | 4 – 6 hours |
| Before date of conference call | Familiarise yourself with information provided for a few species in order to contribute to discussion for these species in particular on conference call | 1 - 2 hours |
| Date of conference call (tbc) | Conference call to discuss and revise scores | 4 hours |
| Date of conference call (tbc) | **Round 2** - Rescoring | This can be completed during the call |
| Between conference call and 17 January | Email round 2 scores back to Rike, ideally immediately after the call | 1 minute |

**Data Protection and confidentiality**

You will be asked for demographic information, which, along with your probability scores, will be kept in password-protected spreadsheets on a computer at Newcastle University, UK. Your name will be disaggregated from your answers.

The results of the expert elicitation exercise (median and agreement between scores) will be published in a peer-reviewed journal, but individual scores will not be identifiable to person.

**Authorship**

If you agree to participate and complete both the scoring and take part in the conference call, you will be offered co-authorship of the scientific publication of this work. We intend to submit the paper to Conservation Letters.

**Will participation prejudice me in any way?**

Your participation in this study is completely voluntary. Should you wish to withdraw at any stage, or to withdraw any comments that you have supplied, you are free to do so without prejudice.

**Where can I get further information?**

This research has been granted approval by the Newcastle University Ethics Committee (Reference 15388/2018). Should you require any further information, or have any concerns, please do not hesitate to contact Dr Rike Bolam or Professor Phil McGowan.

#### Instructions

**Timeline**

See Table 1.

**Rules**

Please do not talk to the other evaluators during scoring for Round 1 OR Round 2. You can use any other means available to answer the questions - e.g. talk to outsiders, consult literature, draw on past experience, acquire and interpret data. It would also be appropriate for you to draw on your own knowledge and experiences of regions or species.

**Information and data entry**

We will make information available to you for every candidate species. You will also be able to use any other information you have access to. The information provided by us will include the following for all species:

- The population estimate and direction of trends for 1993, 2010, and 2019, as far as known
- Generation lengths
- Past and current threats to the species
- Conservation actions that have taken place
- A justification for why the conservation actions the species received in one or both periods are considered plausibly sufficient to have prevented its extinction, given the magnitude of threats and the species' population and distribution

We will send this information on an excel spreadsheet, alongside the questions, so you can answer the questions directly in the spreadsheet. We have also uploaded the species information online, in case you prefer reading the species information in a different format. It is available here.

**Questions**

We will ask you the same questions for all species. These questions are:

1. Realistically, what do you think is the **lowest plausible probability** that, if action had ceased in 1993, and no subsequent actions were implemented, the species would have gone extinct by 2020? [Answers from 0 – 100, e.g. a score of 70 means that there is a 70% probability that the species would have gone extinct by 2020 if action had ceased in 1993 and no actions had been implemented after that year]
2. Realistically, what do you think is the **highest plausible probability** that, if action had ceased in 1993, and no subsequent actions were implemented, the species would have gone extinct by 2020? [Answers from 0 – 100, e.g. a score of 70 means that there is a 70% probability that the species would have gone extinct by 2020 if action had ceased in 1993 and no actions had been implemented after that year]
3. Realistically, what is your **best estimate for the probability** that, if action had ceased in 1993, and no subsequent actions were implemented, the species would have gone extinct by 2020? [Answers from 0 – 100, e.g. a score of 70 means that there is a 70% probability that the species would have gone extinct by 2020 if action had ceased in 1993 and no actions had been implemented after that year]
4. Additionally, if species met the criteria to be included for the time period 2010 – 2020, the following questions were also asked: Realistically, what do you think is the **lowest plausible probability** that, if action had ceased in 2010, and no subsequent actions were implemented, the species would have gone extinct by 2020)? [Answers from 0 – 100, e.g. a score of 70 means that there is a 70% probability that the species would have gone extinct by 2020 if action had ceased in 2010 and no actions had been implemented after that year]
5. Realistically, what do you think is the **highest plausible probability** that, if action had ceased in 2010, and no subsequent actions were implemented, the species would have gone extinct by 2020)? [Answers from 0 – 100, e.g. a score of 70 means that there is a 70% probability that the species would have gone extinct by 2020 if action had ceased in 2010 and no actions had been implemented after that year]
6. Realistically, what is your **best estimate for the probability** that, if action had ceased in 2010, and no subsequent actions were implemented, the species would have gone extinct by 2020? [Answers from 0 – 100, e.g. a score of 70 means that there is a 70% probability that the species would have gone extinct by 2020 if action had ceased in 2010 and no actions had been implemented after that year]

In other words, if you give a probability score of 0%, then you are saying that the species would have persisted even without conservation action, for the time period under consideration. On the other hand, if you give a probability score of 100%, then you are saying that the species would not have persisted without conservation action, and would have gone extinct for the time period under consideration.

#### Frequently asked questions

**What makes someone an expert?**

For this study we believe that you have sufficient knowledge to help make an estimate with regards to the questions. Good expert performance is about:

- Having a holistic understanding of the subject matter
- Always seeking the truth
- Knowing the limitations of your knowledge
- Producing success when practicing your expertise

We want you to have a go at every question. Our question format will enable you to communicate your uncertainty for each question.

**The questions are impossible!**

We have tried to make the questions as clear as possible by only asking about two data points. However, there is always variability, particularly in natural systems. This is why we ask you to communicate your uncertainty to us by communicating a realistic upper and lower bound that would capture this uncertainty. We then ask you to think about what the most likely outcome will be and communicate this to us as your best guess.

**How did we identify candidate species?**

To qualify as candidates, the species:

- Must be likely to be extant currently
- Must have had a significant threat or suite of threats that might have driven it extinct in the absence of actions. This included that the species would have had a small population, or substantial declines, during one or both of the time periods.
- Must have received some significant conservation actions during the period. Conservation actions must have had, or be likely to have had, a positive impact on the species, i.e. either led to an increase or slowed a decline of the species.

'Conservation action' encompasses the full range of interventions, including protected area establishment and management, legislation (e.g. to prohibit hunting), control or management of invasive alien species, control of hunting/trapping, habitat restoration, and species recovery interventions such as captive breeding, translocation, supplementary feeding, nest-site provision etc.

The screening of species was done by two separate groups. Mammals were identified by Louise Mair as well as Marco Angelico at the Global Mammal Assessment. The results were then compared, and any species with disagreement were discussed to reach consensus on inclusion of the species.

**How do we define extinction?**

For the scores, we would like to know whether the species would have gone **extinct in the wild** if not for conservation action. We also mean that the last individual in the wild would have disappeared by 2020, and so do not mean functionally extinct. In the paper, we will also list those species that are currently listed as Extinct in the Wild on the IUCN Red List, to ensure they are included.

**What do the probability values reflect?**

We have identified a range of categories that the probability values correspond to, see Table 2.

*Table 2. Range of probabilities and their meaning for whether extinction was prevented through conservation action (adapted from Keith et al., 2017).*

| **Range of probabilities** | *Was extinction prevented through conservation actions?* |
|---|---|
| 0.99 - 1.00 | The actions are **virtually certain** to have prevented extinction, i.e. would have prevented extinction in 99 of 100 species similar to the target. There is a less than one in a hundred chance that the taxon would have persisted without conservation action during the period. |
| 0.90 - 0.98 | The actions are **very likely** to have prevented extinction, i.e. would have prevented extinction in 49 of 50 to 19 of 20 similar species. There is a one in fifty to one in twenty chance that the taxon would have persisted without conservation action during the period. |
| 0.75 - 0.90 | The actions are **quite likely** to have prevented extinction, i.e. would have prevented extinction in 19 of 20 to three in four similar species. There is a one in twenty to one in four chance that the taxon would have persisted without conservation action during the period. |
| 0.50 - 0.74 | The actions are **more likely than not** to have prevented extinction, i.e. would have prevented extinction of three-quarters of similar species. There is a one in four to 50:50 chance that the taxon would have persisted without conservation action during the period. |
| 0.25 - 0.49 | The actions are **quite possible but unlikely** to have prevented extinction, i.e. would have prevented extinction in one quarter to one half of similar species. There is more than a 50:50 and up to a 3 in 4 chance that the taxon would have persisted without conservation action during the period. |
| 0.10 - 0.24 | The actions are **quite unlikely** to have prevented extinction, i.e. would have prevented extinction of one tenth to one quarter of similar species. There is a 3 in 4 to 9 in 10 chance that the taxon would have persisted without conservation action during the period. |
| 0 - 0.09 | The actions are **very unlikely** to have prevented extinction, i.e. would have prevented extinction of up to one tenth of similar species. There is more than a 9 in 10 chance that the taxon would have persisted without conservation action during the period. |

**How do we define when an extinction has been prevented?**

We will measure agreement amongst the evaluators for all questions, but the best estimate will be used as an indication of extinctions prevented. We will summarise the overall results in terms of the number of species whose extinction has been prevented as X-Y, with X representing species with a median best estimate >50% (i.e. more likely than not to have had their extinction prevented) and Y representing species with a median best estimate >90% (i.e. very likely) to have had their extinction prevented), following an analogous approach for defining Possibly Extinct and Extinct species (Butchart et al., 2018).

We have also visualised the probabilities and when extinction was prevented, to help you with your assessment (Fig. 1).

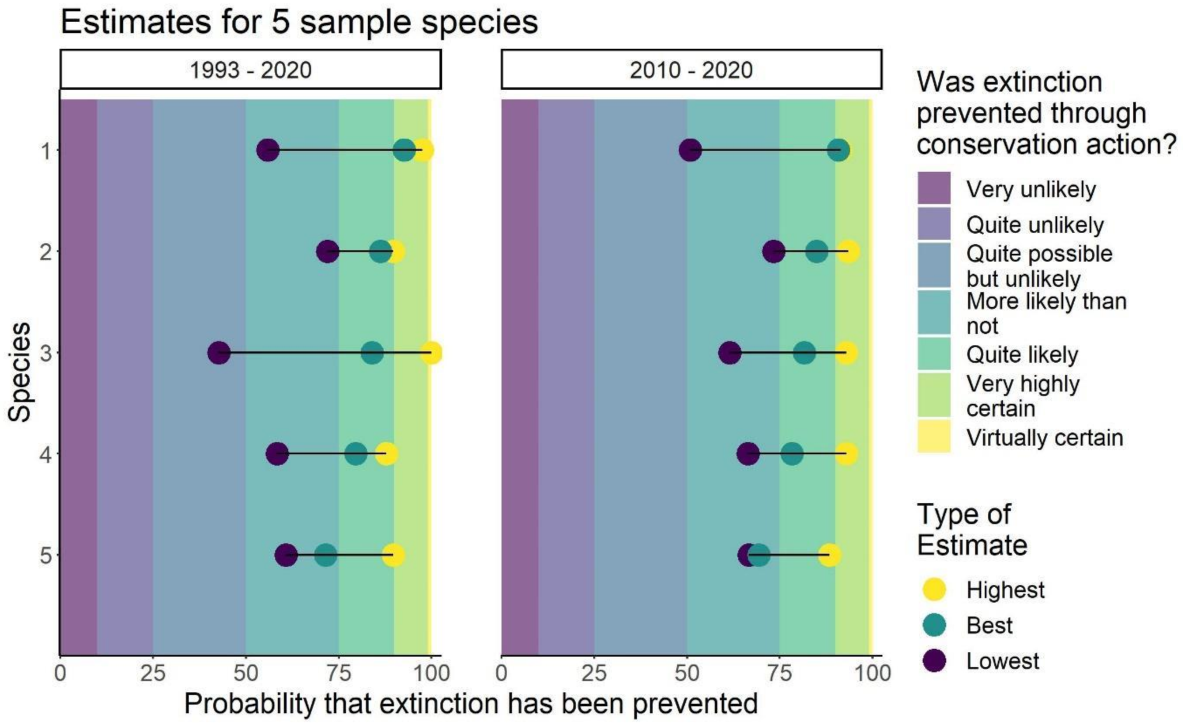


*Figure 1. Example of highest, best and lowest estimate for probabilities for five species.*
